# Chromosome 3p deletion leads to extensive genomic alterations in diverse cancers and confers synthetic lethality in uveal melanoma

**DOI:** 10.64898/2026.01.21.700826

**Authors:** Mitchell C. Cutler, Porter B. Howland, Miroslav Hejna, Jun S. Song

**Affiliations:** Department of Physics, University of Illinois Urbana-Champaign, Urbana, IL, USA; Carl R. Woese Institute for Genomic Biology, University of Illinois Urbana-Champaign, Urbana, IL, USA; Cancer Center at Illinois, University of Illinois Urbana-Champaign, Urbana, IL, USA

**Author notes:** Correspondence (J.S.S.). These authors contributed equally (Mitchell C. Cutler, Porter B. Howland).

**Keywords:** Uveal melanoma, isochromosomes, fragmentation of chromosomes, cancer genome evolution, synthetic lethality

## Abstract

Chromosome 3p (chr3p) is frequently deleted in multiple cancers, indicating the presence of shared tumor suppressors. Analysis of genomic alterations in 33 different cancer types implicates the deletion or deleterious mutations of SET-domain-containing 2 (*SETD2*) at chr3p21 in significantly facilitating the formation of isochromosomes, consisting of two identical mirror-imaged arms, thereby promoting genomic instability conducive to large-scale chromosomal rearrangements and rapid cancer genome evolution. Fracturing of dicentric isochromosomes during cell division is pervasive and follows the dynamic fragmentation pattern of solids under impulse. Across cancers, isochromosomes form most frequently on chr8 to amplify the *MYC*-containing q-arm and chr17 to delete the *TP53*-containing p-arm. In the most aggressive uveal melanoma (UVM) subtype, chr3 deletion also includes *MITF*, a critical melanocyte differentiation and survival factor, and co-occurs with chr8q amplification. We demonstrate that MITF is a master transcriptional regulator of *GNAQ*/*GNA11*, mutated in 90% of UVM patients, and the associated synthetic-lethal genes identified by recent CRISPR screening studies. *MITF* maintains MAPK and calcium homeostasis in UVM, and its deletion is thus accidental, creating an early crisis during oncogenesis. We further show that MITF, MYC, and GNAQ/GNA11 form coupled regulatory feedback loops in the melanocyte lineage, and *MITF* deletion in UVM creates acute dependency on MYC-mediated rescue via chr8q amplification, often as a consequence of isochromosome formation. The discovered feedback loops predict both overall and relapse-free patient survival within the most aggressive UVM subtype, explain sensitivity to therapeutic gene perturbations, and inform effective combinatorial therapies.

## INTRODUCTION

Uveal melanoma (UVM), arising from the uveal tract melanocytes, is the most common primary intraocular cancer in adults, second to skin cutaneous melanoma (SKCM) in frequency among all melanoma cases in the US^1^. Co-occurring chr3 deletion and chr8q amplification (8q+) found in ∼50% of UVM patients^2,3^ comprise the most aggressive subtype with poor relapse-free survival and elevated metastasis risk^2,4^. Chr3 monosomy (M3) is thought to be an early oncogenic alteration in the evolution of aggressive UVM, likely preceding^5^ the accompanying 8q+, and partly reflects the need to delete the tumor suppressor gene *BAP1*^6^, the second retained copy of which is often mutated to completely inactivate the gene^2,3,7^. Consistent with this putative ordering of events, recent single-cell sequencing^8^ of a primary tumor with high metastatic potential showed the branching of ancestral M3 clones into large subclones subsequently acquiring 8q+. These phenomena thus suggest a strong selection pressure on cancer cells to amplify 8q in response to M3, but the source of this selection pressure and associated therapeutic opportunities remain unknown. In addition, over 90% of UVM patients harbor mutually exclusive mutations in the G-protein α-subunit genes *GNAQ* and *GNA11*. We show that hidden within these molecular stratifications resides hitherto-unrecognized information about the trajectory of cancer evolution and the selection pressure that forced the UVM cells to depend on a melanocyte-specific regulatory network.

The 8q+ event in M3 UVM often involves isochromosomes, consisting of two identical mirror-imaged arms and simultaneous deletion of the complementary arms (Figure 1A). Upon close examination of 10,632 patient datasets from 33 different The Cancer Genome Atlas (TCGA) cancers^3^, we have found isochromosomes to be linked to chr3p (3p) deletion and contribute to chromosomal instability (CIN) across multiple cancers. CIN is a cellular state in which large-scale chromosomal rearrangements can emerge from mis-segregation of chromosomes during cell division^9^. CIN is thought to occur in bursts, especially at early stages of tumor evolution^10^, and is a major source of rapid genome diversification on which natural selection can act during oncogenesis. Recent studies have identified multiple roles of SETD2 in protecting normal cells against CIN. First, SETD2 methylates α-tubulin in microtubules during mitosis, a step required for proper mitotic spindle formation, and the loss of this function leads to genomic instability^11,12^. Second, SETD2 is the sole enzyme that can catalyze the trimethylation of H3K36 (H3K36me3)^13^, and the loss of H3K36me3 in SETD2-deficient cells promotes the formation of isochromosomes^14^; isochromosomes can function as a chromatin bridge between dividing cells and initiate CIN^14–16^. Finally, SETD2 loss causes nuclear morphological defects, promoting aneuploidy and CIN^17^. Even though the essential role of SETD2 in preventing isochromosomes has been demonstrated in cell lines^14^, the impact of its aberration in cancer patients has not yet been demonstrated. Of note, *SETD2* is located at 3p21 and thus deleted in M3 UVM. Strikingly, in UVM, isochromosomes are not observed on chr3, but frequently occur on chromosomes 1, 6, and 8, leading to 1p-/1q+, 6p+/6q-, and 8p-/8q+ (Figure 1B); furthermore, 8q+ isochromosomes occur only in the context of M3 in TCGA-UVM, suggesting that *SETD2* loss likely precedes and facilitates 8q+ in M3 patients. Extending this observation, our analysis of pan-TCGA cancers implicates *SETD2* loss in forming isochromosomes in diverse human cancers. Across cancers, isochromosomes occur most frequently on chr8, amplifying *MYC* as in UVM, and on chr17, deleting *TP53*. We further show that dicentric isochromosomes are prone to fragmentation, which can be accurately modeled using dynamic fracture analysis^18,19^, and that such repeated fragmentation can cause ultra-high amplification of oncogenes.

**Figure 1.**
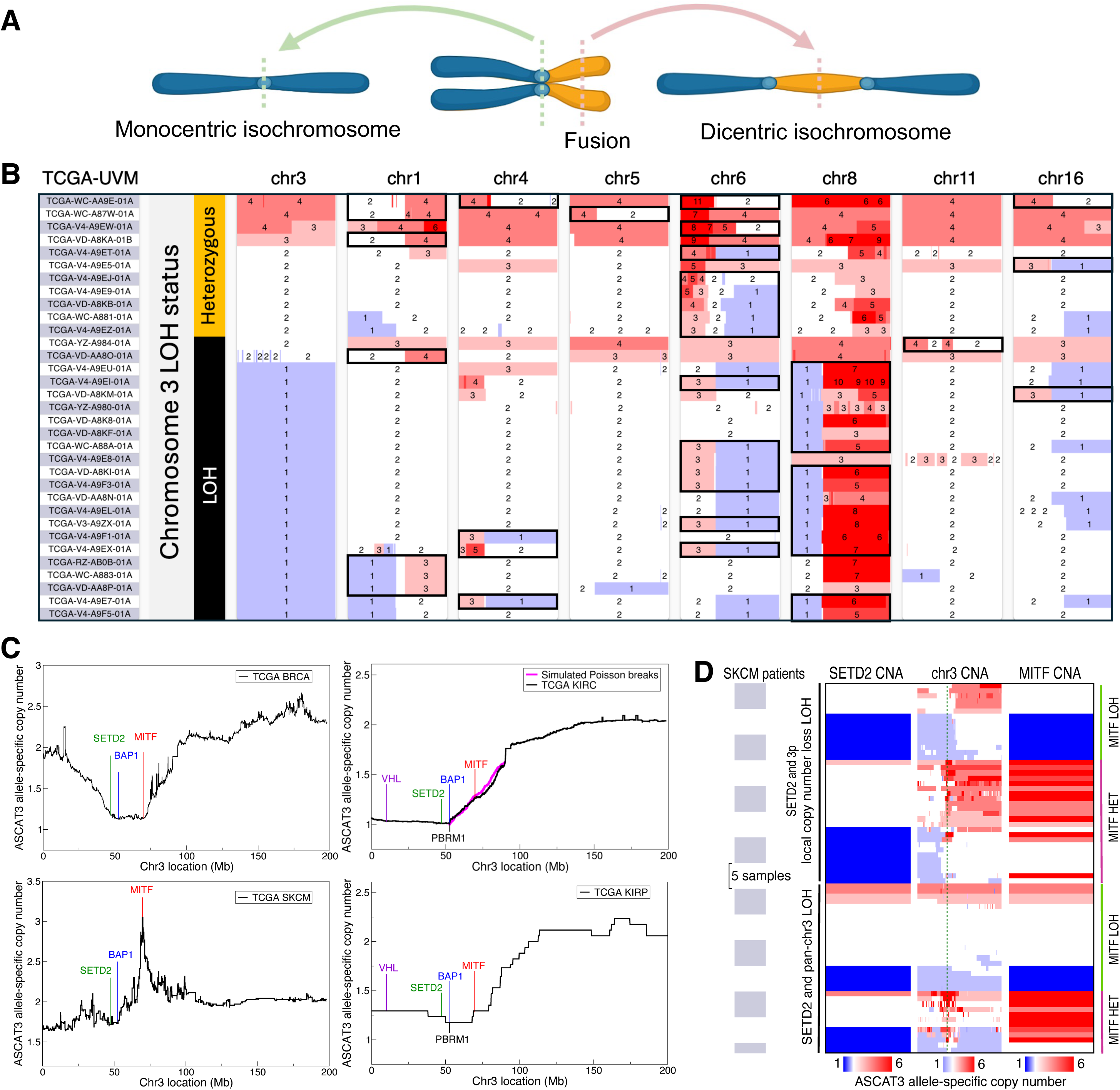
Chr3 deletion precedes isochromosome formation in UVM and is linked to deep MITF amplification in SKCM. (A) Illustration of monocentric and dicentric isochromosome formation. (B) ASCAT3 allele-specific copy number data for TCGA-UVM patients with isochromosomes (black boxes). Isochromosomes are significantly enriched in the group with chr3 LOH (Binomial test p-value = 2.9 × 10^−4^ ). Isochromosomes on chr8 occur only in the context of M3 (Fisher’s exact test p-value=3.0 × 10^−7^). (C) Mean chr3 deletion profile in four TCGA cancers (142 BRCA, 338 KIRC, 70 SKCM, 17 KIRP patients). The KIRC (top right) plot shows a fitted Poisson break model (magenta line) initiated at the centromere to the location of *BAP1*. (D) Copy number alterations (CNA) showing loss (blue) and gain (red) in SKCM patients with either partial or full LOH on chr3. Green dashed line in the middle “chr3 CNA” panel indicates *MITF*. All focal *MITF* amplification loci in the context of chr3 LOH are heterozygous (HET). (B) and (D) are modified versions of UCSC Xena browser screenshots^85^.

Chr3 loss in UVM embodies another hidden driving force. The *GNAQ*/*11* mutations induce constitutive activation of the G-proteins G_q_ and G_11_, leading to the sustained production of diacylglycerol (DAG) and inositol 1,4,5 trisphosphate (IP3). Elevated DAG activates RAS-dependent MAPK pathway by binding and activating the guanine nucleotide exchange factor RASGRP3^20^, creating critical dependence of *GNAQ/11*-mutant cells on RASGRP3. Furthermore, recent high-throughput screening assays have identified synthetic-lethal genes that accompany the *GNAQ*/*11* mutations^21^. In particular, constitutively synthesized IP3, if not degraded via the phosphatase INPP5A, can accumulate and increase intracellular calcium ions to a toxic level. Targeting RASGRP3 or INPP5A with small molecules in the context of *GNAQ*/*11* mutations thus represents a promising strategy for treating a large fraction of UVM patients^21^. However, the UVM dependency on RASGRP3 and INPP5A in the background of M3 poses a perplexing paradox.

Namely, we show that both *RASGRP3* and *INPP5A*, as well as several other critical UVM-vulnerability genes^21,22^, are bound by and transcriptionally activated by MITF, the master regulator of melanocyte differentiation and survival^23^. Despite being a critical hub in the network of synthetic-lethal genes, however, *MITF* resides at 3p14 and thus gets co-deleted with *SETD2*/*BAP1* in M3 UVM. Our discovered regulatory network thus predicts that acute MITF depletion reduces UVM-vulnerability genes and creates lethality in *GNAQ*/*11*-mutant UVM, as supported by independent screening experiments^22,24^, implying that *MITF* deletion is an accident that M3 UVM cells must resolve. We here show that MITF and MYC regulate each other’s transcription and form a positive feedback loop in melanocytes and that the M3-mediated crisis of *MITF* deletion poses a strong selection pressure on UVM to amplify *MYC* on 8q24 to restore the pre-M3 level of MITF. The positive feedback loop between MITF and MYC thus explains the coupling of M3 and 8q+ and resolves the paradox. This newly gained insight, in turn, uncovers an opportunity to develop combinatorial synthetic-lethal therapeutics that can benefit UVM patients.

## RESULTS

### The pattern of 3p deletion reflects shared selection pressure to delete *SETD2* in multiple cancers

Chr3, in particular the p arm, is often lost in diverse cancers, including breast cancers^25^ (∼30%), renal carcinomas^26,27^ (∼60%), pancreatic adenocarcinoma^28,29^ (PAAD) (∼20%), and UVM^3^ (∼50%). In UVM, entire chr3 is lost, making the search for potential tumor suppressor genes difficult (Figure S1A). We thus sought to learn from the pattern of 3p deletion in TCGA breast cancer (BRCA), clear-cell renal cell carcinoma (KIRC), papillary renal cell carcinoma (KIRP), skin cutaneous melanoma (SKCM), and PAAD data. Aligning the copy number profiles of only those patients who had a deletion event on 3p revealed similarities and key differences (Figure 1C, Figure S1B; Methods). Notably, the 5.2 Mb region containing *SETD2* and *BAP1* was preferentially deleted in all five cancers; but, *MITF* deletion was prevalent only in BRCA and KIRP, neutral in KIRC (as simulated by Poisson breaks), under negative selection in SKCM (Figure 1C), and mostly neutral in PAAD, but with one patient who had deep *MITF* amplification in the background of *SETD2*/*BAP1* deletion (Figure S1B). These results strongly supported that in addition to the known tumor suppressor BAP1, SETD2 inactivation also was targeted by multiple cancer types.

### *MITF* amplification often arises from either 3p or pan-chr3 deletion in SKCM

*MITF* is amplified in ∼10% of primary and ∼20% of metastatic SKCM, leading to the lineage-addiction hypothesis proposing that certain cancers depend on lineage-specific pathways that are regulated by critical differentiation factors and promote cancer initiation/progression^30^. In TCGA-SKCM data, however, 17 (3.7%) and 30 (6.5%) out of 461 patients had a single copy of *MITF* and *SETD2*, respectively; 28 patients who had *BAP1* deletion also had *SETD2* deletion (Figure 1C); and, 6 of the 30 patients with *SETD2* copy=1 had focal amplification of *MITF* (Figure 1C,D). This pattern indicated that, as was the case in UVM, *BAP1*/*SETD2* deletion facilitated skin cutaneous melanomagenesis, but the reduced deletion frequency and focal amplification of *MITF* in the background of *BAP1*/*SETD2* deletions also suggested that *MITF* deletion in the melanocyte lineage might have been an undesirable accidental consequence of the proximity between the three genes. Closer investigation revealed that the *BAP1*/*SETD2* locus actually harbored higher frequency of cryptic deletions: 39 patients (8.5%) had local copy-number-loss (CNL) loss of heterozygosity (LOH) of *SETD2* and 33 patients (7.2%) had pan-chr3 LOH, sometimes interspersed by many heterozygous fragments (Figure 1D). Importantly, 20 (28.8%) of these 72 patients had focal *MITF* amplifications, all in the heterozygous *MITF*-locus context (Figure 1D); furthermore, 42 patients had duplicated the intact chr3, likely at a later stage of cancer progression after the initial *BAP1*/*SETD2* deletions had already facilitated oncogenesis. Co-occurring partial deletion of 3p and focal amplification of *MITF* could be explained by break-fusion-bridge^31^ cycles of dicentric chromosome 3, while focal *MITF* amplification in the background of pan-chr3 LOH possessed signatures of extensive fragmentation in chromothripsis^31,32^ (Figure 1D). Together, these results strongly supported that a large fraction of *MITF* amplification in SKCM was associated with *BAP1*/*SETD2* deletion.

We thus partitioned the *MITF* amplification events into 4 groups: shallow amplification (copy=3,4) with or without *SETD2* LOH, and deep amplification (copy≥5) with or without *SETD2* LOH. For shallow amplification, copy number alteration (CNA) in the presence of *SETD2* LOH (8%) exhibited a propensity to selectively amplify *MITF*; in the absence of *SETD2* LOH (92%), however, there was no evidence of specifically amplifying *MITF* over other genes on chr3 (Figure S1C). Deep amplifications showed a clear signature of positive selection on focal amplification of *MITF*, with 19 (34.5%) out of the 55 deep amplifications occurring in the background context of *SETD2* LOH and showing ∼2-fold higher amplification than when chr3p was heterozygous (Figure 1D, Figure S1C; Table S1A). Deep *MITF* amplifications in SKCM were thus significantly associated with *SETD2* LOH (Fisher’s exact test p-value = 4.7 × 10^−6^, odds ratio = 6.3), while shallow *MITF* amplifications did not exhibit a trace of selection specifically for *MITF* itself, unless 3p was partially deleted. These results thus revealed that partial 3p deletion crucially contributed to amplifying the oncogene *MITF* in SKCM and supported that *MITF* deletion in UVM was an accidental byproduct of entire chr3 loss.

### Deleterious mutations of *SETD2* are significantly associated with isochromosome formation in papillary renal-cell carcinoma

Frequent *SETD2* mutations were previously reported in the TCGA-KIRP cohort^27^. Given the uncertainty in determining the functional consequence of missense mutations, we focused on unambiguous deleterious mutations. We identified 9 KIRP patients harboring a stop-codon gain or a frame-shift mutation in *SETD2* at a minimum allele-frequency of 0.5. Out of these 9, 5 patients had 3 ± 2 isochromosomes (Table S1B). We found a significant association between the high-frequency deleterious *SETD2* mutation status and the presence of isochromosomes in 269 KIRP patients with intersecting data (Fisher’s exact test p-value = 3.0 × 10^−4^, odds ratio = 16.4). Other TCGA cancers, including KIRC, did not have enough patients with high-frequency deleterious *SETD2* mutations to test for association.

### Deletion of *SETD2* or *PSIP1* is strongly associated with isochromosome formation in the majority of TCGA cancer types

In addition to deleterious point mutations, cancer cells often suppress a gene’s function by deleting the gene. Importantly, a critical gene deletion event, e.g. transiently activating CIN, might later get compensated by amplifying the second copy to avoid mitotic catastrophe^33^. We thus examined not only copy number changes but also the LOH status of *SETD2* in 33 TCGA cancer types, spanning 10,632 patients (Methods). In addition to *SETD2* on chr3, the only known writer of H3K36me3, we also examined *PSIP1* on chr9, the only known reader of H3K36me3 involved in homologous recombination and RAD51 recruitment to double-strand breaks (DSBs)^34^. The LOH status of *SETD2* was significantly associated with isochromosome formation in non-acrocentric chromosomes in 10 (30%) out of 33 TCGA cancers (Wald-test FDR threshold of 10^−2^, Figure 2A; Table S1C,D; Methods); chr3 was excluded in this analysis to avoid correlations arising from over-counting isochromosomes that themselves led to *SETD2* LOH. Likewise, *PSIP1* LOH was significantly associated with isochromosome occurrence, excluding chr9, in 17 (52%) out of 33 TCGA cancers (Wald-test FDR threshold of 10^−2^, Figure 2A), sharing all 10 cancers with the *SETD2* LOH result. *SETD2* LOH and *PSIP1* LOH synergized, and the double-LOH status of *SETD2*/*PSIP1* was significantly associated with isochromosome occurrence compared to the complement set and the double-heterozygous *SETD2*/*PSIP1* cohort, with the most pronounced effect seen in KIRP (Figure 2A, Table S1D). Isochromosomes occurred most frequently on chr8 (99% 8q+ including *MYC*, 1% 8q-), followed by chr3 (96% 3p-including *SETD2*/*BAP1*, 4% 3p+) and chr17 (98% 17p-including *TP53*, 2% 17p+) (Figure 2B). Isochromosomes thus led to critical oncogene amplifications and tumor suppressor deletions across cancer types.

**Figure 2.**
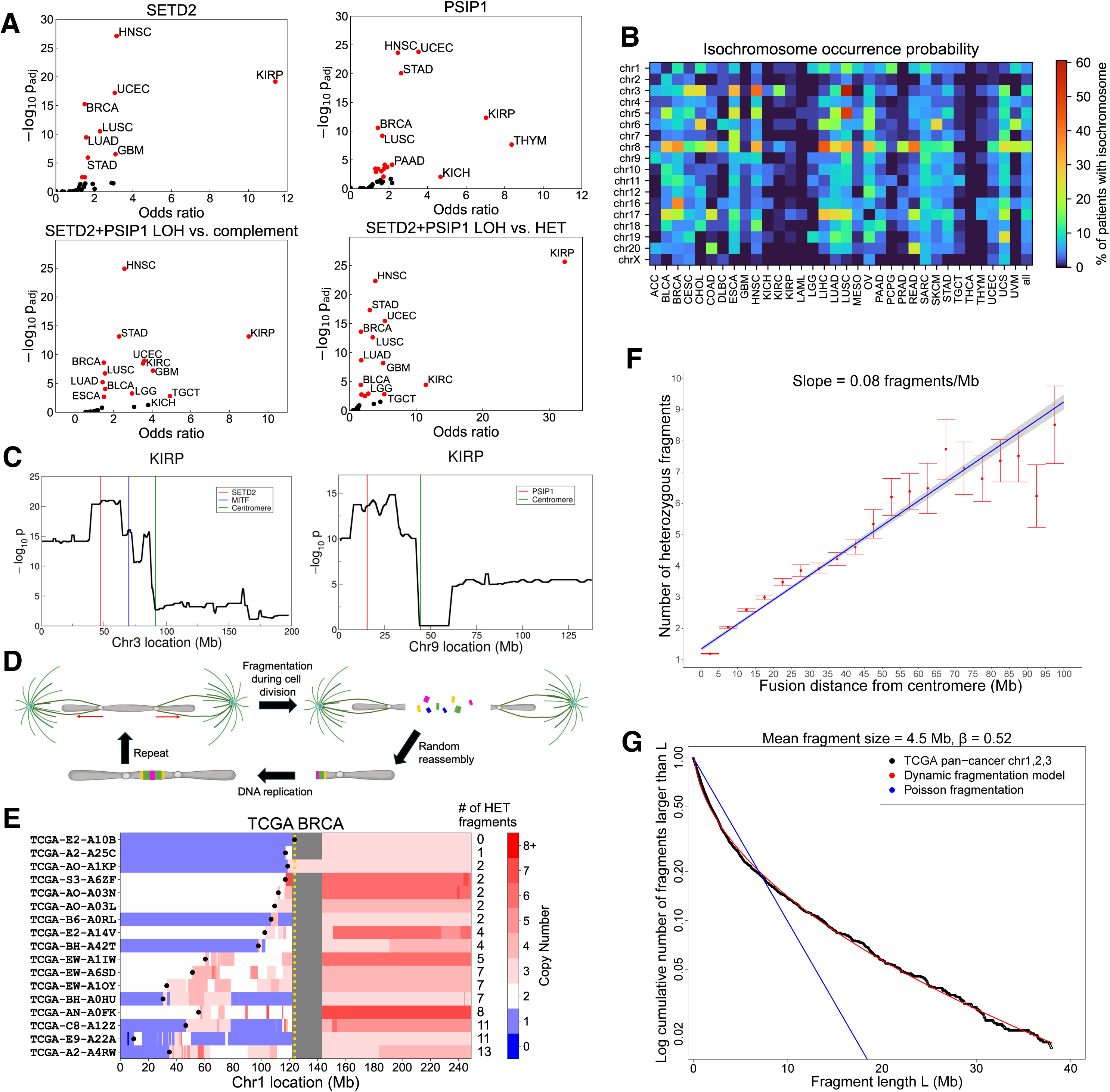
*SETD2* and *PSIP1* deletions are associated with isochromosome formation and chromosome fragmentation across multiple cancers. (A) Benjamini-Hochberg-corrected Wald-test p-values comparing isochromosome formation rates in patients with vs. without LOH in either *SETD2* (top left) or *PSIP1* (top right), and in patients with double-LOH *SETD2*/*PSIP1* vs. the complement (bottom left) or double-heterozygous *SETD2*/*PSIP1* (bottom right). Red indicates passing the statistical significance threshold (*p_adj_* < 0.01) for each of the 33 TCGA cancers. (B) Isochromosome rates as probability of occurrence per patient for each cancer type; e.g., ∼50% of LUSC patients have isochromosome 5. (C) Wald-test p-value for observing higher isochromosome occurrence probability in patients with LOH at each chromosome location compared to patients without LOH at the same location in TCGA-KIRP. Chr3 (left) and chr9 (right) locations were evenly sampled at 100kb intervals, and the transformed p-values were smoothed using a running window of 20 for visualization. (D) Illustration of how a dicentric isochromosome can fragment during cell division. (E) Examples of increased fragmentation as a function of the distance from centromere (yellow dotted line) to fusion point (black dot) on chr1 isochromosomes of TCGA-BRCA patients. Gray regions indicate unavailable data. (F) Number of heterozygous fragments on the LOH arm as a function of the distance of fusion site from centromere on 11,658 isochromosomes in 33 TCGA cancers. The shaded region indicates the 95% confidence interval of fragment numbers predicted by univariate linear regression against the distance, including an intercept. (G) Fragment-size distribution of 2188 isochromosomes detected on metacentric chr1,2,3 in 33 TCGA cancers. The patient data (black) agree well with a dynamic fragmentation model (red) and deviates from the Poisson model using the same estimated mean (blue).

As segmental deletions and amplifications tend to have long-range correlations, we further examined the statistical association of LOH status at specific locations on entire chr3 or chr9 with isochromosome occurrence on the remaining chromosomes. We performed the analysis in 7 TCGA cancer types that showed the highest levels of association between *SETD2*/*PSIP1* LOH and isochromosome formation (Figure 2C, Figure S2A,B). In 6 out of the 7 cancers, the 3p arm containing *SETD2* showed much higher association than the 3q arm with isochromosome formation on the remaining non-acrocentric chromosomes; KIRC was an exception to this pattern, because we removed chr3 isochromosomes which were most prevalent in that cancer. In 5 out of the 7 cancer types, the 9p arm containing *PSIP1* clearly showed much higher association with isochromosome formation than the 9q arm; in head-and-neck squamous cell carcinoma (HNSC), even though the p arm was more significant than the q arm, the LOH status on the q arm itself showed very high association with isochromosomes, reflecting the fact that 185 HNSC patients (36%) had pan-chr9 LOH. BRCA and UCEC showed elevated association on the q arm, suggesting that there might be another gene on chr9q contributing to isochromosomes.

### Dicentric isochromosomes show massive fragmentation between centromeres

Isochromosomes can be either monocentric, if fused at the centromere, or dicentric, if fused at the boundary of partial arm-deletion^14^ (Figure 1A). As a dicentric chromosome can form a bridge between dividing cells and fragment^31^ (Figure 2D), we counted the number of heterozygous copy-number fragments on the arm experiencing LOH. The number grew linearly as a function of the distance between fusion site and centromere, yielding 1 new fragment every ∼12Mb on average (Methods; Figure 2E,F, Figure S2C). Dicentric isochromosomes were also significantly more fragmented than non-isochromosomes (Mann-Whitney U-statistic p-value< 5 × 10^−324^, 1.9-fold enrichment; Methods). Furthermore, the size distributions of heterozygous fragments between centromeres on dicentric isochromosomes showed excellent fitting to dynamic fragmentation models of solids under transient force^18^ (Figure 2G, Figure S2D; Methods). The fitted exponent 𝛽 being less than 1 indicated heterogeneity in fracture propensity across each chromosome and deviation from homogeneous Poisson fragmentation (Figure 2G). These results thus supported that isochromosomes served as a major source of complex genome alterations, yielding numerous fragmented and amplified segments, reminiscent of chromothripsis^31,32,35^. Some of these amplified fragments contained oncogenes related to cancer progression and poor patient survival. For example, ultra-deep amplifications of the driver genes *CCND1*^36^ and *GAB2*^37^ in TCGA-BRCA arose from the fragmentation of chr11q-isochromosomes (Figure S2E,F), similar to the amplification mechanism of *MITF* in SKCM (Figure 1C,D, Figure S1C).

### MITF is a central hub in UVM cancer codependency network

A recent CRISPR-screening study by Elbatsh *et al*.^21^ reported 29 UVM synthetic-lethal genes, the therapeutic depletion of which significantly impeded proliferation in the presence of *GNAQ*/*11* mutations. To understand the relation among these hit genes, we examined their top 100 codependency genes from the Cancer Dependency Map (DepMap)^24^ database and found that *MITF* was the top shared gene. Examining the screening data showed that the effect of *MITF* knockout (KO) correlated strongly with the reported hits, but perhaps did not meet the threshold criteria employed in that study (Figure S3A). Of note, another independent screening study found *MITF* to be among the top 10 UVM-specific essential genes^22^. We thus included *MITF* in the hit list and constructed a codependency graph for the 30 genes (Figure 3A); the highest betweenness centrality score of *MITF* confirmed the visible role of *MITF* as a central hub in the network (Figure 3B). This central role of *MITF* was consistent with the strongly selective effect of its perturbation on melanocyte-lineage cancers in DepMap screening experiments (Figure S3B). The effect of *MITF* KO was most acute in UVM, even more pronounced than in SKCM. These results thus highlighted the prominent role of MITF in UVM biology and strongly supported that *MITF* deletion in the M3 subtype was accidental, as was the case in SKCM (Figure 1C,D, Figure S1C, Figure S3B)

**Figure 3.**
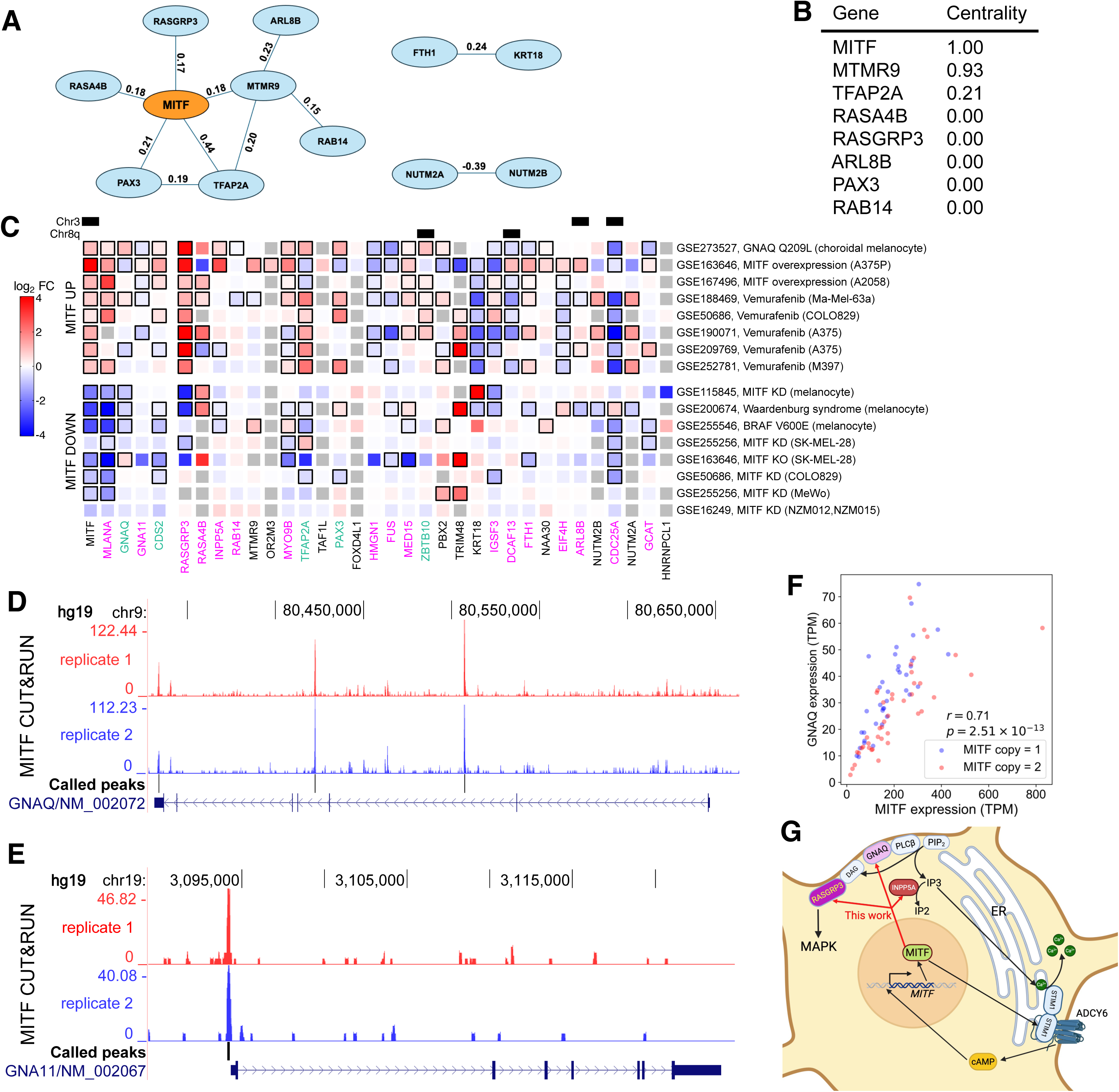
MITF is a master regulator of the network of synthetic-lethal genes in UVM. (A) A network of UVM synthetic-lethal genes and MITF constructed from DepMap’s top 100 codependency genes and corresponding correlation scores^24^. Genes with degree zero are not shown. (B) The relative betweenness centrality scores for the 8 genes in the first connected component of the codependency network in (A). (C) Heatmap of log_2_ fold-change (FC) of expression levels in MITF overexpression (top block) and MITF knockdown (KD)/KO/mutation (bottom block) experiments in melanocyte-lineage cells. The figure shows differential regulation of the vulnerability gene CDS2^22^, 29 synthetic-lethal genes from Elbatsh *et al*.^21^, and *GNAQ*/*11*. The pigmentation gene *MLANA*, a known transcriptional target of MITF, is included as a positive control. Black bordering boxes indicate statistically significant differential expression (adjusted p-value < 0.05). Missing data are shown in gray. MITF binding in promoter and intron regions, marked by magenta and teal gene names, respectively, support direct regulation of the indicated genes by MITF. (D) MITF CUT&RUN data in the melanoma cell line SK-MEL-28^38^ showing binding in *GNAQ* introns. (E) Same as (D), but for *GNA11* promoter. (F) Scatter plot of *MITF* and *GNAQ* expression levels in TCGA-UVM. Pearson correlation coefficient 𝑟 is calculated by pooling the M3 and D3 patients together. (G) Proposed regulatory network of MITF and GNAQ/11 in UVM.

### MITF is a direct transcriptional regulator of synthetic-lethal genes and forms a double-positive regulatory feedback loop with mutated GNAQ/11

To further elucidate the role of MITF in melanocytes, we examined published experimental data modulating MITF expression and found that MITF transcriptionally regulated many of Elbatsh *et al*.’s 29 synthetic-lethal genes^21^, as well as cytidine diphosphate diacylglycerol synthase 2 (CDS2) identified by Chan *et al*.^22^ (Figure 3C). In particular, *RASGRP3*, the downstream effector of GNAQ/11 mutations and the key mediator of the MAPK pathway in UVM^20^, was consistently elevated upon MITF overexpression; conversely, MITF suppression consistently reduced *RASGRP3* expression (Figure 3C). Furthermore, MITF cleavage under targets and release using nucleosome (CUT&RUN) data^38^ in melanocyte-lineage cells showed direct binding of MITF in the *RASGRP3* promoter (Figure S3C). The top synthetic-lethal target^21^ *INPP5A* and the UVM-vulnerability gene^22^ *CDS2* similarly exhibited overall activation by MITF across multiple independent experiments (Figure 3C) and showed direct binding of MITF in their promoters (Figure S3C). Strikingly, among the 29 synthetic-lethal targets, 15 were significantly differentially expressed in 5 or more of the 8 MITF-overexpression experiments; likewise, 15 synthetic-lethal targets showed significant differential expression in 2 or more of the 8 MITF-suppression experiments, with only 7 showing no statistically significant change. Furthermore, MITF CUT&RUN data^38^ showed that 16 synthetic-lethal genes were bound by MITF within 1kb of the transcription start site (TSS), with additional four genes bound by MITF in introns (Table S1E). Together, these results demonstrated that MITF directly regulated a large subset of the synthetic-lethal genes in UVM.

Given MITF’s regulation of multiple genes on which UVM critically depended, we next investigated whether MITF also directly regulated *GNAQ*/*11*, the two most frequently mutated genes driving UVM progression. MITF CUT&RUN data^38^ showed binding in *GNAQ* introns and *GNA11* promoter (Figure 3D,E); the expression levels of *MITF* and *GNAQ*, and also *GNA11* to a lesser extent, were highly correlated in UVM (Figure 3F, Figure S3D). MITF perturbation data^39^ in human primary melanocytes expressing either wild-type BRAF or BRAF^V600E^ confirmed the transcriptional regulation of many synthetic-lethal genes by MITF, with *GNAQ* again showing stronger correlation than *GNA11* (Figure S3E). In turn, the expression levels of *MITF* itself and the synthetic-lethal genes regulated by MITF were significantly elevated in choroidal melanocytes engineered^40^ to express GNAQ^Q209L^ (Figure 3C). The increased expression level of *MITF* may be attributed to the phosphorylation of its regulator CREB by Ca^+2^-dependent protein kinases^41^ or Ca^+2^-activated protein kinase A (PKA)^42^. Alternatively, the recently discovered coupling of Ca^+2^ sensing and cAMP generation^43^ could also contribute to the elevation of *MITF* expression in *GNAQ/11*-mutant UVM (Figure 3G, Discussion). These results thus showed that MITF and GNAQ/11 formed a double-positive regulatory feedback loop.

Together, these findings thus explained why MITF was a central hub in the dependency network (Figure 3A), but also raised a paradox that MITF, the master regulator of the synthetic-lethal genes, was lost during cancer evolution in ∼50% of UVM patients, potentially causing a crisis that cancer cells had to surmount. This paradox also posed an accompanying puzzle: why was MITF not a top hit in Elbatsh *et al*.’s CRISPR screening, despite being a central hub? We shall now resolve both enigmas by explicating the transcriptional interplay of MITF and MYC in the context of M3 and 8q+.

### MITF and MYC form a transcriptional regulatory feedback loop in melanocytes

Mean CNA in TCGA-UVM showed frequent global loss of chr3 and whole-arm 8q+ (Figure 4A,B), and these alterations were linked in individual patients (Figure S4A,B), as previously observed^2,3^. Surprisingly, *MITF* and most of its known transcriptional targets were not differentially expressed between M3 and disomy 3 (D3) genotypes (Figure 4C, Figure S4C), indicating that 8q+ might compensate for *MITF* deletion. One potential mechanism was MYC on 8q24, known to rescue MITF’s general transcriptional activity in melanocytes in the absence of MITF^44^. Consistent with this hypothesis, *MYC* copy number was significantly increased in patients with *MITF* deletion (Figure 4D); however, *MYC* expression level again did not correlate with its copy number (Figure 4E). The lack of correlation between expression and copy number for both MITF and MYC indicated the presence of a confounding factor, such as a transcriptional feedback loop for maintaining homeostasis.

**Figure 4.**
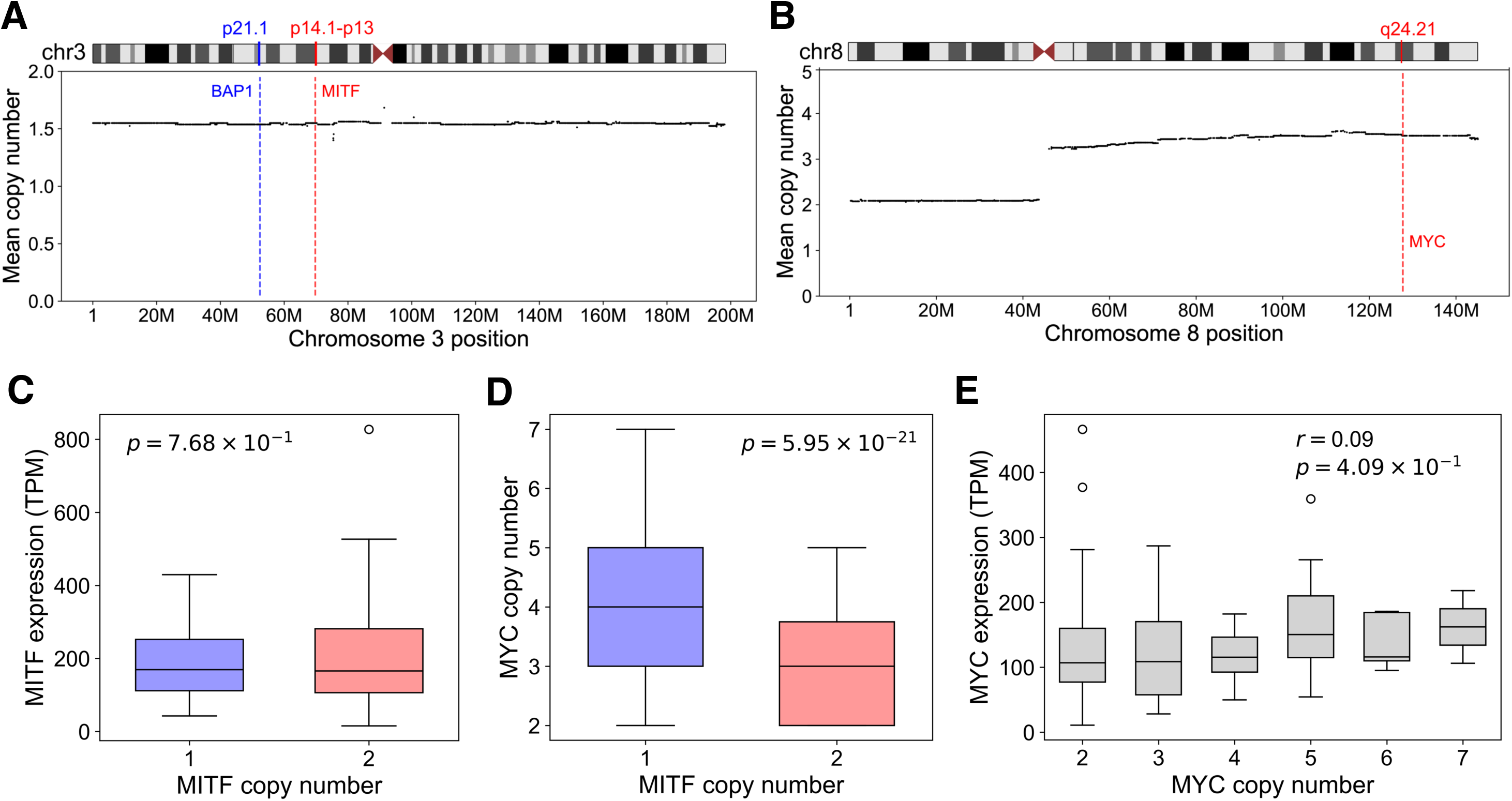
Copy number alterations alone cannot explain the expression levels of MITF and MYC in TCGA-UVM. (A) Averaged pattern of chr3 deletions in TCGA-UVM showing that entire chr3 tends to be lost. ABSOLUTE gene-level copy number data are shown. (B) Same as (A), but for chr8 amplifications, showing that only the q arm tends to be amplified. (C) Boxplots of *MITF* expression levels in M3 vs. D3 patients. Expression level is seen to be sustained when one copy of *MITF* is lost (Wilcoxon rank-sum test). (D) Boxplots of *MYC* copy numbers in M3 vs. D3 patients. *MYC* amplification is significantly associated with *MITF* deletion (Wilcoxon rank-sum test). (E) Boxplots of *MYC* expression levels grouped into distinct copy numbers, showing the lack of Pearson correlation.

Independent ChIP-seq^45^ in melanocyte and CUT&RUN^38^ in melanoma revealed that MITF bound the promoter of *MYC* as well as *PVT1*, the adjacent long non-coding RNA known to regulate MYC expression^46,47^ (Figure 5A); conversely, MYC:MAX bound *MITF* in melanocyte^48^ and 13 different ENCODE^49^ cell lines (Figure 5B, Table S1F). Consistent with these findings, *MITF* and *MYC* expression levels showed strong correlation in UVM (Figure 5C). Indirect regulation of *MYC* by MITF was also previously reported, with MITF depletion leading to significant suppression of MYC^44^. Moreover, several independent MITF modulation experiments confirmed its transcriptional activation of MYC (Figure 5D). In particular, *MITF* and *MYC* were both significantly elevated in choroidal melanocytes expressing^40^ GNAQ^Q209L^ (Figure 5D).

**Figure 5.**
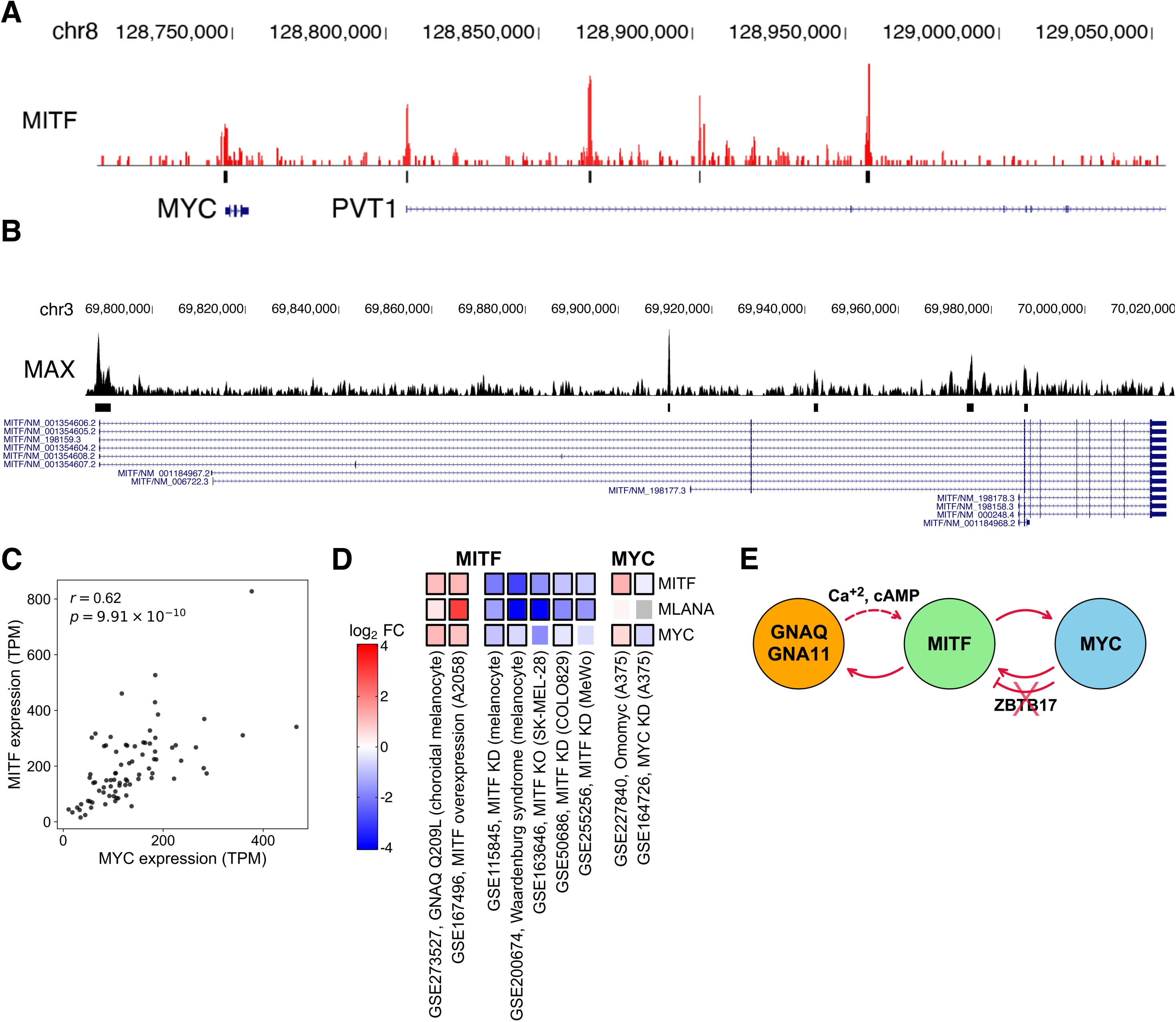
MITF forms coupled double-positive regulatory feedback loops with MYC and GNAQ/11 in UVM. (A) MITF ChIP-seq^45^ in melanocyte, showing binding in the promoter of *MYC* and *PVT1*. (B) MAX ChIP-seq^48^ in melanocyte, as a proxy for MYC, shows binding in the promoters of several *MITF* isoforms. (C) Scatter plot of *MITF* and *MYC* expression levels in TCGA-UVM, showing a strong Pearson correlation. (D) Heatmap of log_2_-FC of expression levels of *MITF*, *MYC* and *MLANA* in MITF overexpression and KD/KO/mutation experiments in melanocyte-lineage cells. MITF and MYC are seen to regulate each other, forming a double-positive feedback loop. *MLANA* serves as a positive control. Black boxes indicate statistical significance (adjusted p-value < 0.05). (E) Coupled gene regulatory network of GNAQ/GNA11, MITF and MYC in UVM. Depletion of ZBTB17 in UVM switches MYC:MAX into an activator of *MITF*.

Regulation of *MITF* by MYC was more complex. To assess whether *MITF* level changed together with other known targets of MYC, we computed the distribution of correlations between *MITF* and other coding genes in UVM. Compared to background, MYC targets showed significantly greater correlation with *MITF* (Figure S5A,B). However, although MYC KD in A375 suppressed *MITF* in one study^50^, treatment of A375 with the MYC-inhibitor Omomyc increased the level of *MITF*^51^ (Figure 5D), informing that MYC could either activate or repress *MITF* in a context-dependent manner. Supporting this idea, it is known that although the MYC:MAX dimer functions as an activator, ZBTB17 (a.k.a. MIZ1) can interact with MYC and MYC:MAX to turn them into repressors, and this switch depends on the ratio of MYC and ZBTB17 binding^52^. We found evidence that ZBTB17 bound^49^ several locations within *MITF* (Figure S5C), and its expression was anti-correlated with *MITF* in SKCM (Spearman 𝜌 = −0.27, p-value= 2.7 × 10^−&^), but not in UVM (Spearman 𝜌 = 0.27, p-value= 0.02). Importantly, in TCGA-UVM, *ZBTB17* located on chr1p was preferentially co-deleted with *MITF* (Binomial test p-value=0.012), sometimes because of isochromosome (Figure 1B), and deleted preferentially in 8q+ patients (Binomial test p-value= 5.6 × 10^−5^). These findings thus explained the previous report that genes typically repressed by the MYC:MAX:ZBTB17 complex were selectively activated in 8q+ patients^3^ and supported the transcriptional activation of *MITF* by MYC. As ubiquitination and subsequent degradation of ZBTB17 via HUWE1 could also help switch MYC activity from repression to activation^53^, we examined the effect of *HUWE1* on *MITF* expression. Bivariate regression of *MITF* expression against *MYC* and *HUWE1* expression showed significant partial correlation with both genes (Figure S5D). These results thus demonstrated that MYC activated *MITF* transcription in UVM. In summary, we discovered coupled double-positive feedback loops involving GNAQ/11, MITF and MYC (Figure 5E), which could function together to maintain UVM homeostasis.

### *MYC* amplification rescues UVM from accidental *MITF* deletion

Given the aforementioned regulatory feedback, we next examined the effect of *MYC* amplification on *MITF* expression. Whereas *MITF* expression diminished upon *MITF* deletion with *MYC* copy=2, *MYC* amplification restored the expression to D3 levels, with no significant difference between *MITF* copy=1 with *MYC* amplification and *MITF* copy=2 (Figure 6A). Conditioned on *MITF* deletion, *MITF*, *RASGRP3* and *INPP5A* all showed statistically significant elevation with *MYC* amplification (Figure 6B-D), restoring or even slightly increasing the levels of *RASGPR3* and *INPP5A* relative to the D3 condition (Figure 6E-F). *MYC* expression could also be predicted by *MYC* amplification, conditioned on *MITF* copy=1 (Figure S5E), supporting that the negligible marginal correlation was confounded by the regulatory feedback between MITF and MYC (Figure 4E). These results strongly supported that *MYC* amplification effectively restored the expression levels of MITF and its key target genes critical for UVM oncogenesis. Our findings also informed that the accidental deletion of *MITF* in M3 UVM was initially sensed by MYC through its diminished activation by MITF^44^, leading to 8q+ and 1p-as compensatory mechanisms to maintain the feedback-mediated homeostasis of UVM (Figure 5E).

**Figure 6.**
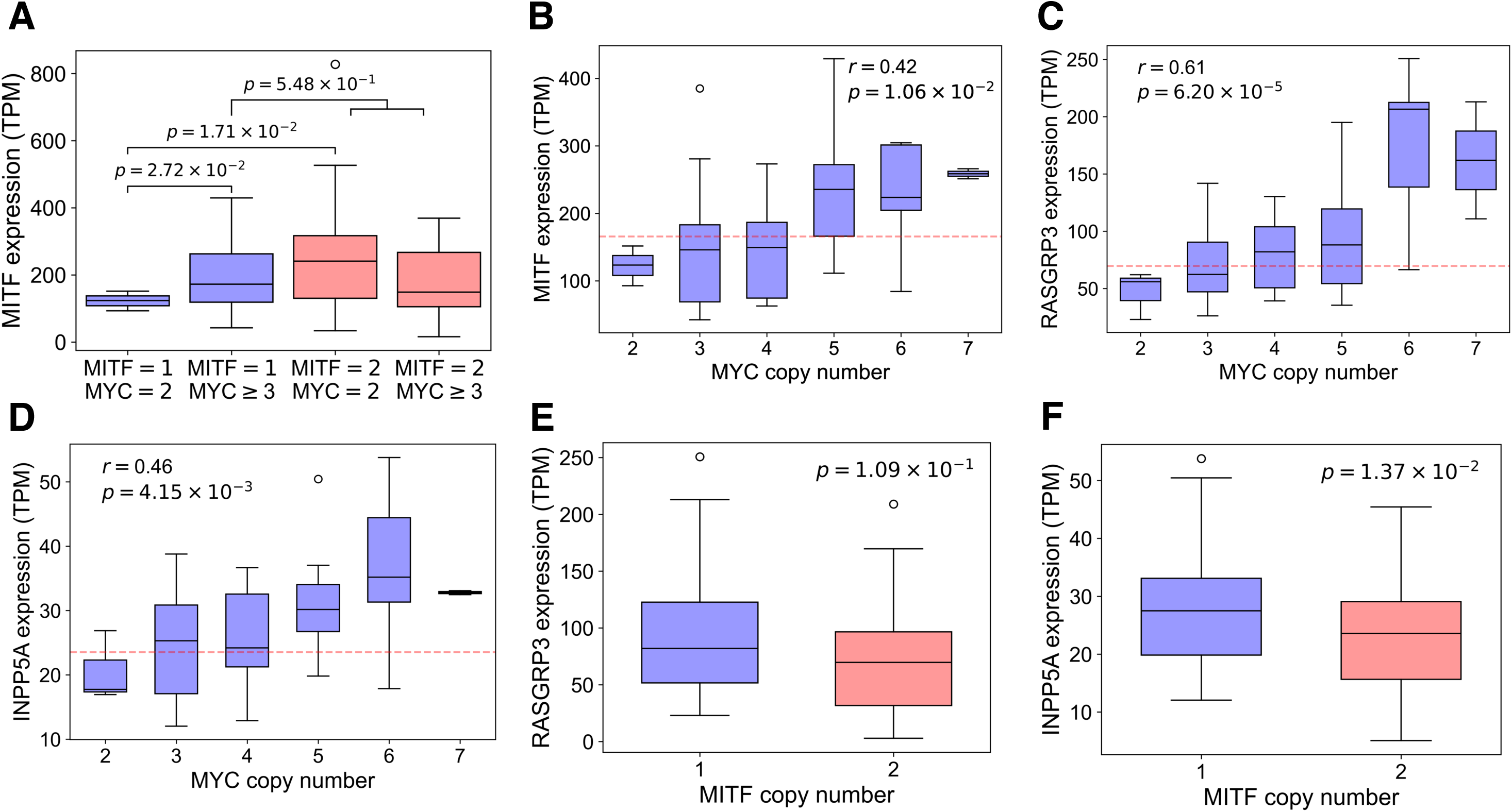
The regulatory feedback loop between MITF and MYC restores the expression of *MITF* and synthetic-lethal genes in M3 patients. (A) Distributions of *MITF* expression levels in patients grouped by *MITF* and *MYC* copy numbers. Single-copy *MITF* with wildtype (WT) copy number *MYC* results in significantly lower expression relative to either single-copy *MITF* with amplified *MYC* or WT copy number *MITF*. However, *MYC* amplification effectively compensates for *MITF* deletion. The t-test p-values are shown. (B) Conditioned on *MITF* deletion, *MITF* expression level can be explained by *MYC* copy number, supporting that *MYC* amplification rescues *MITF* expression in M3 patients (Pearson 𝑟 = 0.42). Dotted red line indicates median *MITF* expression in the D3 group. (C,D) Same as (B), but for MITF-regulated *RASGRP3* and *INPP5A*, respectively. (E) Boxplots of *RASGRP3* expression levels in M3 vs. D3. The difference between the two groups is not significant (Wilcoxon rank-sum test). (F) Same as (E), but for *INPP5A*. The M3 group had a slightly higher level than D3 (Wilcoxon rank-sum test).

In terms of modeling, synergistic dual positive feedback loops with nonlinear interactions are commonly found in genetic networks to create bistability^54^. Moreover, coupling multiple double-positive feedback loops, similar to the MITF/MYC/GNAQ interactions, was shown to be the ideal network architecture for creating bistable states robust against stochastic fluctuations^55^. The existence of bistable states was supported by the bimodal distributions^56^ of *GNAQ*, *MITF* and *MYC* expression in M3 patients harboring *GNAQ*/*11* mutations, jointly exhibiting a high(GNAQ,MITF,MYC) or a low(GNAQ,MITF,MYC) state (Figure S5F), akin to the MITF^low^ and MITF^high^ states in SKCM^57,58^. By contrast, GNA11 did not fit a simple bimodal distribution and contained an extra low(GNAQ,MITF,MYC)/high(GNA11) state that could not be explained by MITF regulation alone (Figure S5F).

### *MYC* amplification is a significant predictor of survival within M3 UVM patients

UVM patients with co-occurring M3 and 8q+ are known to have poor prognosis^2,3^. Even within the M3 context, we found that TCGA-UVM patients harboring *MYC* amplification exhibited significantly reduced overall survival compared to those with *MYC* copy=2 (log-rank test p-value= 0.018; Figure 7A). In contrast, Johansson *et al.* found no statistically significant relationship between 8q+ and relapse-free survival in UVM patients with M3 (log-rank test p-value= 0.11)^2^. However, aggregating their data with M3 patients in TCGA-UVM showed a significant reduction in relapse-free survival for M3 patients with 8q+, regardless of whether patients with chr3 copy-number-neutral or partial LOH were removed (log-rank test p-value = 0.0095 ; Figure 7B) or retained (log-rank test p-value= 0.011; Figure S6A). These results thus demonstrated that 8q+ synergized with M3 to significantly reduce the survival of M3 patients.

**Figure 7.**
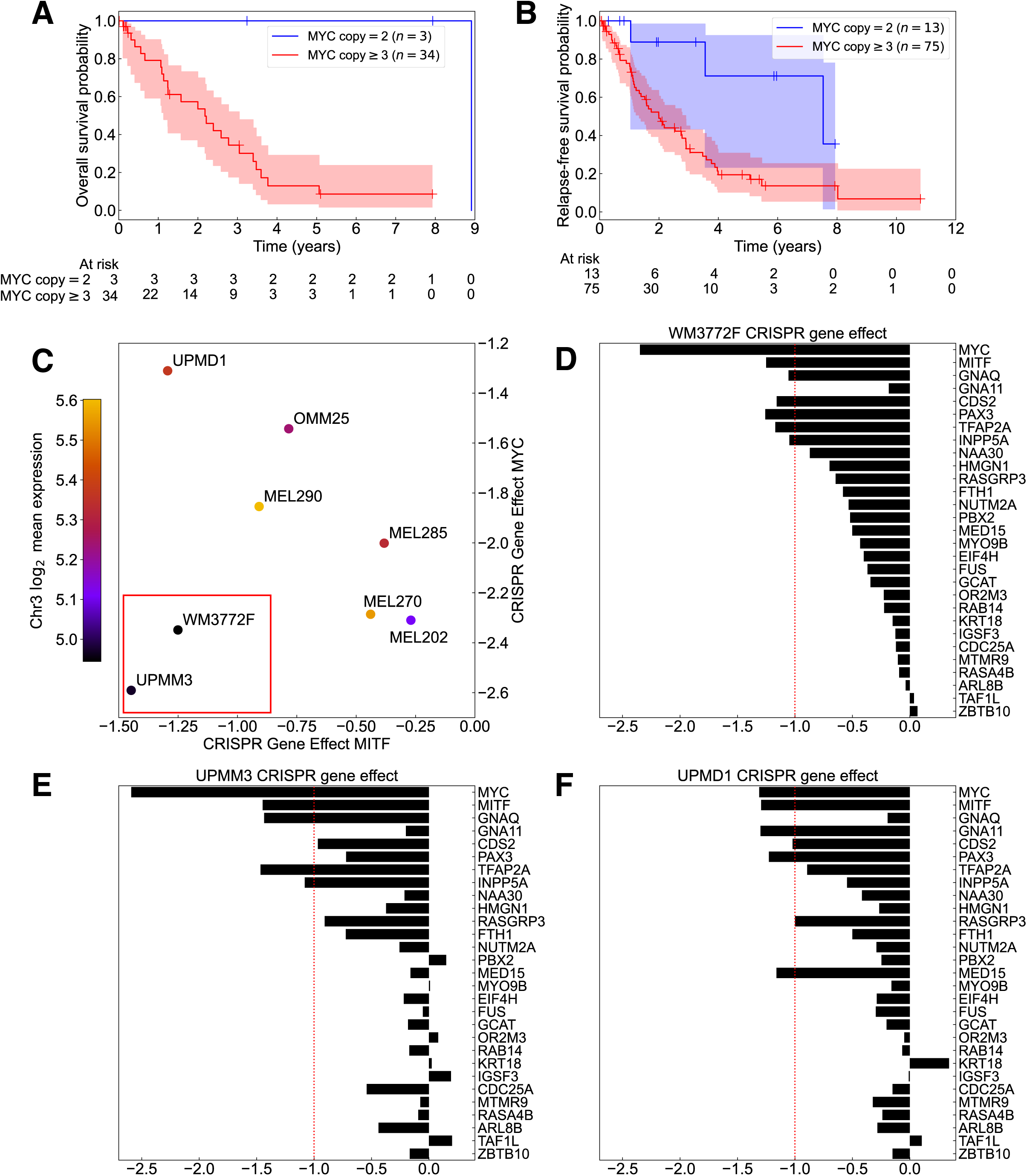
Chr3 deletion worsens prognosis and enhances selective dependence of UVM on MITF and MYC. (A) Kaplan-Meier overall survival curve for TCGA-UVM patients with M3, split into groups with WT or gain in *MYC* copy number (log-rank test p-value= 0.018). The shaded region indicates a 95% confidence interval; crosshairs mark right-censored data. (B) Same as (A), but for relapse-free survival of combined patients from TCGA-UVM and Johansson *et al*.^2^ (log-rank test p-value= 0.0095). (C) DepMap UVM cell lines with available CRISPR gene effect scores for *MITF* and *MYC*. Chr3 log_2_ mean expression is a proxy for chr3 deletion status. UPMM3 and WM3772F (red box) have the lowest chr3 expression and show the highest vulnerability to both *MITF* and *MYC* KO. (D-F) DepMap CRISPR gene effect scores in UVM cell lines for MYC, MITF, GNAQ/11, Elbatsh *et al.*’s^21^ synthetic-lethal genes, and the UVM-vulnerability gene *CDS2*^22^; genes with missing data are excluded. UPMM3 has M3, while UPMD1 has D3. The dashed line at -1 is a recommended threshold for assessing high dependency (lower values).

### M3 enhances the dependency of UVM on MITF, MYC, and synthetic-lethal genes

We reasoned that even though UVM patients with co-occurring M3 and 8q+ had worse prognoses than those with either D3 or D8, they might benefit from a therapy targeting the specific regulatory network on which this UVM subtype critically depended. To identify potential synergistic targets, we examined DepMap’s CRISPR-Cas9 KO gene effect scores^24^ for one UVM cell line with M3/8q+ (UPMM3^59–62^), one with unverified chr3 status but with *MYC* amplification (WM3772F^63^), two with D3/D8 (MEL270^62,64,65^ and MEL290^59,64,66^), and four with D3/8q+ (UPMD1, MEL202, MEL285, and OMM25)^59–61,64,66,67^ (Figure 7C-F, Figure S6B-F).

While all eight UVM cell lines showed strong dependence on *MYC,* a common essential gene, the three with the highest *MITF* dependency were also either *GNAQ*-mutant (WM3772F, Figure 7D; UPMM3, Figure 7E) or *GNA11*-mutant (UPMD1, Figure 7F)^59,60,68^ and depended on either *RASGRP3* or *INPP5A* as well as *CDS2*, consistent with their direct regulation by MITF (Figure 3C, Figure S3C). Two (UPMM3, WM3772F) of these cell lines vulnerable to *MITF* KO had *MYC* amplification and the lowest levels of overall chr3 expression (Figure 7C), with UPMM3 previously verified as being M3^59–62^, and they were strongly dependent on both MYC and MITF (Figure 7D,E). However, UPMD1 possessing D3/8q+, although dependent on MITF, showed the least dependency on MYC among the 8 UVM cell lines (Figure 7C,F). Moreover, *MITF* KO did not have a pronounced effect on other D3 UVM cell lines (Figure S6B-F); likewise, *RASGRP3*, *INPP5A* and *GNAQ* KO also did not have a major effect on two D3 *GNAQ*-mutant cell lines (MEL202, MEL270) (Figure S6B,C). These results thus supported that chr3 inactivation in the context of *GNAQ*/*11* mutations sensitized UVM cells to targeting MITF and MITF-regulated synthetic-lethal genes and elevated their dependency on MYC, needed to compensate for *MITF* suppression (Figure 5E). These enhanced dependencies could be explained by our prediction that an acute therapeutic reduction of MITF or MYC in M3 cells would significantly decrease the levels of GNAQ/11 and synthetic-lethal genes regulated by MITF, recreating the crisis encountered by M3 UVM during early oncogenesis. These findings also shed light on why a CRISPR screening using D3 cell lines might not detect MITF as a top candidate^21^.

## DISCUSSION

Chr3p deletion occurs in multiple cancers^6,12,25,69^ and diseases^70,71^, but understanding the function of this chromosome arm in complex diseases has been challenging. By comparing the pattern of 3p deletion across cancers, we have identified *SETD2* deletion to be shared (Figure 1C) and strongly implicated in forming isochromosomes in human cancers (Figure 2A-C, Figure S2A), as recently proposed^14^. In addition to SETD2, which establishes H3K36me3, the reader PSIP1 of this epigenetic modification is also frequently deleted in patients with isochromosomes (Figure 2C, Figure S2B). Although PSIP1 is the only H3K36me3-binding protein known to be involved in homologous recombination and DSB repair^34^, our analysis suggests that chr9q may also harbor a gene involved in isochromosome formation (Figure S2B). Incidentally, *PHF19* residing on 9q encodes another reader of H3K36me3 which functions as a component of the Polycomb repressive complex 2 (PRC2)^72^. Further investigation is needed to assess whether PHF19 aberration promotes isochromosomes, perhaps by perturbing the transition between open and condensed chromatin structure during replication and chromosome segregation.

This work has analyzed only the copy number LOH information about *SETD2* and *PSIP1* and did not consider their expression levels, because the expression pattern measured post cancer genome evolution may not reflect the relevant level during early oncogenesis. It is, however, plausible that suppression of *SETD2* and *PSIP1* via aberrant transcriptional regulation might also affect their proper function and cause isochromosomes in human cancers^73^ and other diseases^74,75^. In cancer, isochromosomes can have a profound functional impact on genome evolution by creating a transient state of genomic instability responsible for rapid, large-scale genomic rearrangements, likely as an early step during oncogenesis^9,14,31,32,35,76^. By comprehensively analyzing isochromosomes in 33 TCGA cancers, we have shown that dicentric isochromosomes may shatter during cell division, giving rise to massive DNA fragmentation. Despite the complex process of fracture mechanics upon being pulled, the average number of reassembled fragments increases linearly as a function of the distance between fusion point and centromere (Figure 2F), and the size distribution follows the prediction from dynamic fragmentation theory of solids under impulse (Figure 2G, Figure S2D).

This study has also uncovered several hitherto-undocumented critical roles of MITF in both UVM and melanocyte biology. The first role resides in its function as a multifarious regulator of Ca^+2^ homeostasis, required for both GNAQ/11-driven UVM survival and normal melanogenesis. The GNAQ/11-mediated production of IP3 in UVM parallels the pigmentation pathway in normal melanocytes. A well-established function of αMSH that initiates melanogenesis is to bind and activate the melanocortin 1 receptor (MC1R), which predominantly interacts with the G-protein GNAS to generate cAMP, which in turn activates the transcription of *MITF* and other pigmentation-related genes^23^. A less well-understood function of αMSH is to also activate the phospholipase Cβ (PLCβ)^77^ which hydrolyzes phosphatidylinositol 4,5-bisphosphate (PIP_2_) into DAG and IP3 (Figure 3G) – the same two messengers that are constitutively produced in GNAQ/11-driven UVM; it is currently unknown which melanocortin receptor directly interacts with GNAQ and/or GNA11 to initiate this signaling cascade. In normal melanocytes, the elevated level of IP3 then induces Ca^+2^ release from the endoplasmic reticulum (ER) and oligomerization of STIM1 at the ER-plasma membrane junction, which in turn leads to Ca^+2^ influx into melanocytes from the environment and activates the adenylyl cyclase ADCY6 to maintain the generation of cAMP^43^ (Figure 3G). This positive feedback loop between cAMP and Ca^+2^ sensing is thought to be essential for sustaining the high level of cAMP and transport of L-phenylalanine into the cell for robust melanogenesis^77,78^. These phenomena highlight the importance of maintaining the balance of Ca^+2^ concentration for normal melanocyte function.

A recent study has shown that MITF, the master transcriptional regulator of melanocyte differentiation and melanogenesis, transcriptionally activates *STIM1*^78^, the recipient of the message from IP3. We here have shown that MITF also regulates the annihilator of the message, *INPP5A*. This MITF-mediated Ca^+2^ homeostasis mechanism built into melanocytes for melanogenesis may create a strong dependency of melanocyte-derived melanomas, such as UVM, on the clearing potential of INPP5A when evolutionary forces select for potent oncogenic mutations, such as the *GNAQ*/*11* mutations. Melanocytes therefore may have been ready to resolve the constitutive IP3 production and Ca^+2^ flux changes arising as an inadvertent byproduct of *GNAQ*/*11* mutations, but this acute dependency also creates vulnerability and a valuable opportunity for therapy. In the absence of INPP5A, constantly accumulating IP3 triggers p53-dependent apoptosis and thus selective synthetic lethality in GNAQ/11-driven UVM^21^. Supporting this vulnerability, calcium channel blockers have shown efficacy in treating UVM^79^. Intriguingly, RASA4B is a Ca^2+^-dependent suppressor of RAS-MAPK pathway, and its mRNA level is seen to be suppressed by MITF in some contexts and elevated in others (Figure 3C). Further investigation is needed to unravel its function as a synthetic-lethal target in UVM^21^.

The second role of MITF our study has discovered is its transcriptional regulation of *GNAQ*/*11*, as well as *RASGRP3* encoding the guanine nucleotide exchange factor activated by DAG, one of the two messengers constitutively produced by the *GNAQ*/*11* mutations. In *GNAQ*/*11*-mutant UVM, RASGRP3 is a critical mediator of MAPK pathway activation^20^ and a synthetic-lethal target^21^. The aforementioned feedback loop of sustaining the cAMP level initiated by constitutive IP3 production implies a positive feedback loop between MITF and mutant *GNAQ*/*11* gene regulation, explaining the dominant dark skin phenotype of *GNAQ*/*11* mutations observed in mice^80^.

Consistent with our model, choroidal melanocytes engineered to express GNAQ^Q209L^ express a significantly higher level of MITF^40^. In turn, our finding that MITF directly regulates *RASGRP3* further explains why *GNAQ*/*11* mutations increase the *RASGRP3* mRNA level, even in SKCM^20^. The positive feedback loop between MITF and *GNAQ*/*11* thus not only helps sustain the level of *GNAQ*/*11* but also elevates the level of the key effector protein RASGRP3.

The third critical role of MITF lies in its central position at the hub of the co-dependency network of previously reported synthetic-lethal genes (Figure 3A). We have shown that the centrality of MITF in the network can be partly attributed to the direct transcriptional regulation of many synthetic-lethal genes by MITF itself (Figure 3C-F, Figure S3C,D). MITF is thus a comprehensive modulator of the synthetic-lethal gene network. These observations predict that targeting MITF will perturb several synthetic-lethal genes simultaneously in UVM, suggesting an efficacious combinatorial therapy of targeting MITF together with other co-dependent genes using small molecules^21,81,82^. For some UVM subtypes, targeting MITF alone may even suffice to profoundly inhibit the proliferation of cancer cells (Figure 7C-F), and our work has shown that these subtypes are characterized by a specific pattern of recurrent genomic alterations.

Namely, we have found a source of evolutionary selection force that can explain the preferential co-occurrence of M3 and 8q+ in UVM. Being at the hub of the synthetic-lethal gene network and directly regulating many of the essential genes, the inadvertent loss of *MITF* in the M3 background is disadvantageous for UVM, requiring a dosage compensation by amplifying its transcriptional regulator *MYC* on 8q and deleting the repressor *ZBTB17* on 1p. *MYC* amplification is sufficient to bring MITF expression back to the D3 level and thereby rescue UVM cells from the inadvertent reduction of synthetic-lethal genes caused by M3 (Figure 6). This recurrent pattern of genomic evolution found in ∼50% of UVM, representing the most aggressive subtype (Figure 7A,B), thus imposes acute co-dependency between the two oncogenes and suggests that reduced-dose combinatorial targeting of MYC and MITF together with RASGRP3, INPP5A and CDS2 should create potent synthetic lethality in patients harboring both M3 and 8q+ (Figure 7C-E), while minimizing toxicity. Although traditionally considered undruggable, new inhibitors targeting MITF, MYC, or MYC:MAX heterodimerization are under active development^81–83^ and will facilitate the exploration of this therapeutic strategy.

Further study is needed to investigate the role of the regulatory feedback loop between MITF and MYC in SKCM. Unlike UVM, SKCM cancers tend to activate the MAPK pathway by mutating BRAF, and MYC activation has been shown to be sufficient and necessary to develop resistance to BRAF inhibition^84^. As BRAF inhibition leads to MITF accumulation and enhances its transcriptional activity (Figure 3C), our finding that MITF binds *MYC*/*PVT1* and activates *MYC* expression helps explain the transcriptional therapeutic response leading to a MYC-mediated resistance mechanism. Consistent with our hypothesis, MITF suppression increases the sensitivity of SKCM to BRAF inhibition^81^.

Our study has thus identified the evolutionary force driving the genomic alteration landscape of ∼50% of UVM. We have shown that the nested double positive feedback loops involving MITF, MYC and GNAQ/11 are paramount to melanocyte function and that this co-dependency provides an integrated conceptual framework for understanding the previously reported synthetic-lethal targets and the key effectors of *GNAQ*/*11* mutations.

## AUTHOR CONTRIBUTIONS

J.S.S. conceived of, designed, and supervised the project. M.C.C. and J.S.S. performed the isochromosome, survival, and DepMap analyses. P.B.H., M.C.C., and J.S.S. performed the co-dependency network and MITF/MYC perturbation analysis. P.B.H., M.H., and J.S.S. performed the integrative TCGA-UVM analysis and the ChIP-seq/CUT&RUN data analysis. P.B.H., M.C.C., and J.S.S. wrote the manuscript with input from M.H.

## ACKNOWLEDGMENTS

We would like to thank Dr. David E. Fisher for helpful discussions. This project was supported in part by NIH R01CA163336 and the Grainger Engineering Breakthroughs Initiative.

## DATA SHARING PLAN

All code and scripts are freely available at https://github.com/jssong-lab/SPLIT-CHR.

## METHODS

### The Human Reference Genome

Unless explicitly stated otherwise, all genomic coordinates in the manuscript are in hg38.

### The Cancer Genome Atlas (TCGA) data

All data from The Cancer Genome Atlas (TCGA) were accessed through the Genomic Data Commons (GDC) portal (v43.0) or through the UCSC XenaBrowser^85^ (GDC v41.0).

### UCSC Screenshots

The cytobands in Figure 4A,B and the ChIP-seq/CUT&RUN tracks in Figure 3D,E, Figure 5A,B, Figure S3C, and Figure S5C were obtained from visualizing the data on the UCSC Genome Browser^86^.

### Mean ASCAT3 allele-specific copy number analysis conditioned on deletions or amplifications

For a given chromosome arm, deletion-aware copy number alignment was performed by using only those patients who have a chromosomal fragment with total ASCAT3 copy number < 2 on the specified chromosome arm. Amplification-aware alignment was similarly performed by using only those patients who have a chromosomal fragment with total ASCAT3 copy number > 2 on the specified chromosome arm. This approach was used to generate Figure 1C, Figure S1B,C, and Figure S2E.

### Simulation of Poisson-distributed chr3p break points

TCGA-KIRC patients tended to have chr3p contiguous partial-arm deletions including the telomere and *BAP1*, indicating that a single chromosomal break event generated the deletions. To simulate a null model of chr3p break between the centromere and *BAP1* locus, we assumed the break point to be Poisson-distributed between 90 Mb and the *BAP1* TSS. The mean ASCAT3 copy number of 338 KIRC patients with a deletion event on chr3p was 1.627 at the location 90 Mb, theoretically representing 212 patients who still retained two copies at this location and 126 M3 patients. We thus simulated 212 Poisson breaks in the indicated region and computed the mean copy number of the cohort consisting of these 212 simulated patients and 126 M3 patients, yielding the magenta simulated result in Figure 1C.

### Isochromosome calling algorithm

We called isochromosomes by calculating the modal copy number (the most frequently occurring copy number) of each chromosomal arm, using allele-specific ASCAT3 copy number data from TCGA. The locations of centromeres were determined from the “centromeres*”* table from UCSC Genome Browser Table Browser (hg38); p and q arms were demarcated by the midpoint of each centromere, which was determined using the smallest and largest genomic coordinates reported for the centromeric sequence start and end, respectively. Isochromosomes were then identified when all the following conditions were met:

- One arm had LOH, which was enforced by requiring that the modal minor copy number be zero.
- The other arm was heterozygous, which was enforced by requiring that the modal minor copy number be greater than zero.
- The modal major copy number of the arm with LOH was at least two less than the total modal copy number of the heterozygous arm.
- If the modal major copy number of the LOH arm was closer to or equidistant to the modal minor copy number of the heterozygous arm compared to the modal major copy number of the heterozygous arm, then we reasoned that the major copy of the heterozygous arm formed an isochromosome and required its copy number to be at least two. Otherwise, we assumed that the minor copy of the heterozygous arm formed an isochromosome and required its copy number to be at least two.
- The allele-specific minor copy number of the telomeric end of the LOH arm had to be zero. This last condition prevented incorrectly calling non-isochromosome-forming mechanisms that retained the telomere while producing partial-arm LOH.

### Analysis of the association between *SETD2*/*PSIP1* LOH and isochromosome formation

We analyzed the effect of *SETD2*/*PSIP1* LOH on isochromosome formation as follows. Isochromosome occurrences were called in each of the 33 TCGA cancers (10,632 total patients) using the previously outlined procedures for 18 of the 24 human chromosomes; the acrocentric chromosomes 13, 14, 15, 21, 22, and Y were removed.

We determined the *SETD2* or *PSIP1* LOH status for each patient based on the gene’s ASCAT3 allele-specific minor copy number being zero. The possible number of isochromosome calls ranged from 0 to 17 for each patient after the exclusion of acrocentric chromosomes and chr3 or chr9 during the analysis using SETD2 or PSIP1, respectively, and was further reduced to 16 when analyzing the effect of their double-LOH status. No patients carried both SETD2 and PSIP1 LOH in LAML, PRAD, and THCA cancers, which were excluded from the analyses involving *SETD2/PSIP1* double-LOH status. The Wald statistic

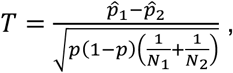

was calculated where 𝑁_1_ and 𝑁_2_ are the number of patients with or without gene LOH, respectively. We calculated the probability of a chromosome being an isochromosome as

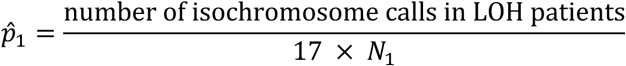

for patients with SETD2/PSIP1 LOH, and similarly for patients with heterozygous SETD2/PSIP1 (substituting 𝑁_1_ for 𝑁_2_ and 𝑝^_1_ for 𝑝^_2_). The probability 𝑝^ is given by

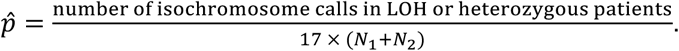

The Wald statistic 𝑇 was evaluated against a standard normal distribution to calculate the p-values for Figure 2A, along with the odds ratio

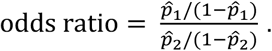

To prevent Type I errors, the FDR was calculated using the Benjamini-Hochberg procedure from SciPy’s (v1.15.1) false_discovery_control and was thresholded at 10^−2^ FDR for calling significance.

In Figure 2C, we computed the Wald statistic comparing the isochromosome occurrence rate in patients with or without LOH at a given location on chr3 or chr9. The locations were sampled every 100kb along chr3 and chr9. For display, we averaged the -log_10_ p-values using a running window of 20 bins.

### Isochromosome occurrence probability calculation

Isochromosomes were called for each of the 33 TCGA cancers for 18 chromosomes (excluding chr13, 14, 15, 21, 22, and Y), using the isochromosome-calling method outlined above. For each cancer type, the number of patients with a specific isochromosome was divided by the total number of patients to calculate the occurrence probability. In the final column (“all”) of Figure 2B, we summed the number of occurrences of each isochromosome across all cancers and divided the sum by the total number of patients (10,632).

### Analysis of chromosomal fragmentation as a function of the distance of isochromosome fusion site from centromere

We analyzed the fragmentation pattern of the isochromosome arm subjected to LOH as follows. The centromere midpoint on each chromosome was determined by first concatenating all HG38 centromeric sequences annotated in the UCSC Genome Browser and then taking the center of the merged region. The p and q arms of each chromosome were then defined to be the left and right side of the corresponding centromere midpoint. Defining heterozygous fragments as the chromosomal fragments in the ASCAT3 allele-specific copy number data with a strictly positive minor copy number, we counted all heterozygous fragments entirely contained in the isochromosome LOH arm. To account for the centromere region itself, we added a count of 1 to the calculated number of heterozygous fragments. The fusion site of isochromosome was determined to be the location of the outer boundary of the farthest heterozygous fragment on the LOH arm from the centromere midpoint. Combining all 11,658 isochromosomes, we used R’s (v4.4.3) linear model (lm; default parameters) to regress the number of heterozygous fragments on the LOH arm against the fusion site distance from the centromere (Figure 2F). Figure 2F (including the fitted line, the error bars, and the 95% confidence interval) was made with ggplot2, using stat_summary_bin for the error bars and geom_smooth for the fitted line and 95% confidence interval. In Figure S2C, the fusion site (“location”), chromosome, and cancer type were all used as independent variables for the linear model fit.

### Analysis of chromosomal fragmentation as a function of dicentric isochromosome status

To compare the number of heterozygous fragments occurring on isochromosomes vs. non-isochromosomes, we called isochromosomes according to the above outlined methods using ASCAT3 allele-specific copy number. Equal copy number fragments immediately flanking the centromere were identified as a single fragment to compensate for missing or incomplete copy number data. The boundaries of the centromere were defined using the smallest and largest genomic coordinates for centromere endpoints listed in the “centromeres*”* table from UCSC Genome Browser. The number of heterozygous fragments were counted in dicentric isochromosomes and non-isochromosomes. Isochromosomes were considered monocentric and excluded from the analysis if they had LOH on an entire arm (divided at the centromere midpoint); otherwise, they were considered dicentric. In cases where the copy number fragment did not span the entire called centromere region, one heterozygous fragment count was subtracted if the copy number of the fragments immediately before and after the centromere region were equal, as this would indicate that the fragments were an artifact of missing copy number data at the centromere. We then tested for the difference between the number of heterozygous fragments in chromosomes with dicentric isochromosome vs. non-isochromosome status using a Mann-Whitney U-test (mannwhitneyu from SciPy v1.15.1; default parameters). The ratio between the mean number of heterozygous fragments with or without isochromosome formation was reported in addition to p-value in the results section “Dicentric isochromosomes show massive fragmentation between centromeres.”

### Mott distribution fitted to the cumulative distribution of chromosomal fragment lengths

Dynamic fragmentation of solids under transient force can be modeled using either stochastic or energy-based physics methods^18^. These theories predict that the cumulative fragment-size distribution decays exponentially in material-dependent power of the fragment size (Figure S2D), and this distribution is called a Mott distribution. The heterozygous fragments on the isochromosome LOH arm were determined as described in the Methods section “Analysis of chromosomal fragmentation as a function of the distance of isochromosome fusion site from centromere.” The intermediate regions with LOH between adjacent heterozygous fragments were discarded, because their fragmentation pattern was lost during cancer evolution. We thus assumed that the observed heterozygous fragments were sampled uniformly from the set of all chromosomal fragments upon fracturing. To demonstrate that the heterozygous fragments occurring on the LOH arm of isochromosomes followed a distribution of dynamic fragmentation predicted by the theory of solids under impulse^18^, we fitted the cumulative number of fragments of length larger than 𝐿 to a Mott distribution,

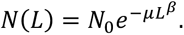

Specifically, the logarithm of the Mott distribution was fitted using R’s (v4.4.3) nonlinear least squares function (nls; default parameters) to numerically determine the parameters 𝜇, 𝛽, and 𝑙𝑜𝑔 𝑁_O_. The mean fragment size of this distribution was calculated to be µ^−1/R^Γ(1 + 1/β). To construct a canonical representative of the fragmentation pattern across multiple cancer types, we aggregated heterozygous fragmentation data from chr1, 2, and 3 (large, metacentric chromosomes) for all 33 cancer types from TCGA (Figure 2G). The aggregated data was then fitted using a Mott distribution (red) and compared to the best-fit Poisson distribution (blue). Because different chromosomes have different arm lengths, we also fitted Mott distributions for individual chromosomes pooled across all cancer types in TCGA (Figure S2D). The LOH status of the acrocentric isochromosomes 13, 14, 15, 21, and 22 could not be reliably determined and were, in addition to chrY, excluded from the analysis.

### Gene network construction and betweenness centrality

The undirected co-dependency network of synthetic-lethal genes (Figure 3A) was constructed with edge weights equal to the Pearson correlation 𝑟 between genes’ CRISPR gene effects (DepMap Public 25Q2+Score, Chronos). The network was restricted to MITF and the 29 synthetic-lethal targets identified by Elbatsch *et al*.^21^, with an edge between two genes included only if either gene appeared in the other’s top 100 co-dependency list. The resulting graph contained three nontrivial connected components (Figure 3A); the remaining vertices, all with degree zero, were not shown. Since only positive edge weights were present, the Pearson correlation edge weights were transformed into edge weight penalties by the formula penalty = 1 − 𝑟, making the analysis of vertex importance amenable to standard techniques based on shortest paths. We then employed the Floyd-Warshall algorithm to compute the shortest paths between all gene pairs and calculate the betweenness centrality score for each vertex in the connected component of MITF. The betweenness centrality of node 𝑣 was computed by

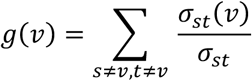

where 𝜎_𝑠𝑡_(𝑣) denotes the number of shortest paths between vertices 𝑠 and 𝑡 which pass through vertex 𝑣, and 𝜎_𝑠𝑡_ denotes the total number of shortest paths between vertices 𝑠 and 𝑡. These centrality scores were then subjected to the normalization

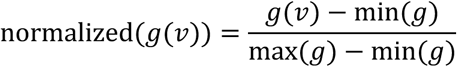

such that the vertex with largest betweenness centrality score was one and the smallest was zero (Figure 3B).

### Summary of MITF or MYC KD/KO/mutation or OE experiments

The curation of multiple OE and KD/KO/mutation data from Gene Expression Omnibus (GEO) for both MITF and MYC followed several standard conventions: if multiple transcript IDs mapped to the same gene name, the data corresponding to the most statistically significant transcript was used. In the case of GSE163646, which contained data from both OE and KO experiments, ENST00000526292.1 was the most statistically significant transcript for MTMR9 during MITF OE but was absent in the MITF KO experiment where instead ENST00000221086 was most significant. To maintain consistency in which isoform’s data was displayed, the MTMR9 log_2_-FC value for GSE163646 MITF KO was removed and instead shown in gray; data for ENST00000526292.1 was shown in GSE163646 MITF OE.

The experiments utilized for the construction of heatmaps are listed below by GEO accession number in top-down order in which they appear in Figure 3C, followed by additional experiments used to corroborate the double-positive feedback loop between MITF and MYC (Figure 5D). Unless otherwise stated, all analyses were performed using GEO’s GEO2R webtool with DESeq2 (v1.49.3)^87^ for differential expression analysis; p-value adjustment was performed within DESeq2 using the Benjamini-Hochberg procedure. GEO Sample accession numbers are provided for each replicate in the groups (case/control) analyzed.

- **GSE273527**^40^utilized choroidal melanocytes cultured from bulbus oculi of human donors. The supplementary materials therein provided differentially expressed genes (DEGs) between RNA-seq of lentiviral transduction-generated mutant GNAQ^Q209L^ vs. normal choroidal melanocytes (12915_2025_2118_MOESM6_ESM.xlsx, sheet name ‘20-RNAseq-NCMvsMutCM DEG’).
- **GSE163646**^38^. MITF OE was accomplished via FLAG tagged MITF in the stable doxycycline inducible overexpression A375P cell line. Differential expression analysis was performed between MITF OE (GSM4983430, GSM4983431, GSM4983432) and empty vector FLAG control (GSM4983427, GSM4983428, GSM4983429); Figure 3C, upper block, second row. Additionally, differential expression analysis was performed for CRISPR-Cas9-generated MITF KO in the human hypotetraploid SK-MEL-28 melanoma cell line (GSM4983417, GSM4983418, GSM4983419, GSM4983420) relative to control (GSM4983413, GSM4983414, GSM4983415, GSM4983416); Figure 3C, lower block, fifth row. Sleuth^88^ (v0.30.2) was used for both MITF OE and KO analyses.
- **GSE167496**. MITF OE in A2058 human melanoma cells was performed in three replicates using lentiviral transduction for MITF OE (GSM5106134, GSM5106135, GSM5106136), and DEGs were computed relative to control (GSM5106131, GSM5106132, GSM5106133).
- **GSE188469**^89^. Differential gene analysis was performed between Vemurafenib-treated replicates (GSM5683486, GSM5683487) relative to a control treated with dimethyl sulfoxide (DMSO) (GSM5683470, GSM5683471) in the metastatic melanoma cell line Ma-Mel-63a harboring the BRAF^V600E^ mutation. Both groups were treated for 24 hours.
- **GSE50686**^45^ treated the COLO829 cell line with the BRAF-inhibitor PLX4032 (a.k.a. Vemurafenib), leading to accumulation of MITF. DEGs were computed between Vemurafenib-treated replicates (GSM1225723, GSM1225724) relative to control (GSM1225721, GSM1225721).
- **GSE190071**^90^. RNA-seq was performed 40 hours post application of the BRAF-inhibitor Vemurafenib (GSM5750555, GSM5750556, GSM5750557, GSM5750558) in the melanoma cell line A375. Three replicates (GSM5750548, GSM5750549, GSM5750550) were used as control.
- **GSE209769**^91^. Replicates of the melanoma A375 cell line were treated with 10 𝜇M of Vemurafenib for 3 days and then sequenced (GSM6395047, GSM6395048, GSM6395049). Differential expression analysis was then performed between the Vemurafenib-treated replicates and control (GSM6395044, GSM6395045, GSM6395046).
- **GSE252781**^92^. The M397 melanoma cell line was sequenced after 12, 24, 36, 48, and 72 hours of BRAF inhibitor (Vemurafenib) application, as well as 12, 24, 48, and 72 hours and 6 days after the removal of Vemurafenib. Controls using DMSO were sequenced at day 0 and 72 and 216 hours after the DMSO treatment. Differential gene expression was performed between replicates treated with Vemurafenib for 12, 24, 48, and 72 hours (respectively: GSM8007258, GSM8007259, GSM8007261, GSM8007262) relative control replicates sequenced at day 0 and after 72 hours of DMSO treatment (respectively: GSM8007257, GSM8345559). Data for the replicate treated with Vemurafenib for 36 hours, the replicate 6 days post Vemurafenib removal, and the control replicate sequenced at 216 hours post DMSO treatment were all unavailable for analysis using GEO2R.
- **GSE115845**^48^. We aligned the raw sequencing reads to the hg38 reference genome using the package HISAT2^93^ (v2.2.1) and assembled transcripts using the package StringTie^94^ (v2.1.1); both used default parameters. We called differentially expressed genes between MITF KD samples (GSM3191783, GSM3191784) and control (GSM3191785, GSM3191786) using DESeq2^87^ (v1.20.0) with default parameters.
- **GSE200674**^95^ utilized melanocytes cultured from induced pluripotent stem cells (iPSCs) derived from a Waardenburg Syndrome (WS) patient carrying a heterozygous mutation in *MITF*. Differential expression analysis was performed between mutant *MITF* (GSM6041871, GSM6041872, GSM6041873) and WT *MITF* (GSM6041874, GSM6041875, GSM6041876) iPSC-derived melanocyte replicates.
- **GSE255546**^39^ used BRAF^V600E^-mutant and WT melanocytes isolated from neonatal foreskin. Differential expression analysis between mutant (GSM8074510, GSM8074511, GSM8074512) and WT (GSM8074504, GSM8074505, GSM8074506) was performed.
- **GSE255256**^96^ used Doxycycline-inducible shMITF KD in MeWo and SK-MEL-28 melanoma cell lines. For MeWo, differential expression analysis was performed between replicates treated with 1 𝜇g/ml Doxycycline (GSM8067677, GSM8067678, GSM8067679) compared to a group without treatment (GSM8067674, GSM8067675, GSM8067676); Figure 3C, lower block, fourth row. Likewise, three SK-MEL-28 replicates were treated with Doxycycline (GSM8067683, GSM8067684, GSM8067685) and compared to control (GSM8067680, GSM8067681, GSM8067682); Figure 3C, lower block, seventh row.
- **GSE50686**^45^ used shMITF in COLO829; differentially expressed genes were computed between shMITF replicates (GSM1225717, GSM1225718) and scrambled controls (GSM1225721, GSM1225722). Differential expression analysis was performed using the R package limma (v3.54.0)^97^, and the microarray platform used was GPL6244.
- **GSE16249** analyzed siMITF KD in two metastatic melanoma cell lines, NZM012 and NZM015. We merged siMITF replicates (GSM409128, GSM409132) in these two cell lines to increase the statistical power of the resulting differential gene expression relative to control (GSM409126, GSM409127, GSM409130, GSM409131). The media control and negative control groups were further merged. The R package limma^97^ (v3.54.0) was used for differential expression analysis. The experiment was done using microarray platform GPL570.
- **GSE227840**^51^ used the MYC inhibitor Omomyc to modulate MYC expression in the BRAF-mutant A375 cell line. Differentially expressed genes were provided in the associated supplementary materials^51^ (‘Supplementary_Tables.xlsx’, sheet name ‘Table S1a A375 Omomyc in vitro’).
- **GSE164726**^50^ performed MYC KD using siRNA in A375. Differentially expressed genes for MYC KD sorted by log_2_-FC (FDR < 0.05) were provided in the associated supplementary materials^50^ (‘Supplementary_Tables.xlsx’, sheet name ‘Supplementary table 2’).

### Construction of signal track figures

MITF CUT&RUN data from Dilshat *et al*.^38^ was acquired from GEO. Called peaks were determined from the associated supplementary materials^38^ (‘elife-63093-supp2.xlsx’, sheet name ‘CutandRunMITFChIPseq-all’). All ChIP-seq and CUT&RUN track figures were made using UCSC Genome Browser^86^.

### Analysis of BRAF^V600E^-mutant melanocytes

Figure S3E was generated using the data GEO GSE255546. We used the ENSG00000136997 Ensembl gene ID for *MYC* when identifying genes in the GEO data. The gene-level counts were normalized by the total counts in each sample and then multiplied by 1000 for visualization.

### Analysis of gene expression levels as a function of *MYC* or *MITF* copy number

For the functional analyses described in Figures 4 and 6, we chose to use the ABSOLUTE copy number data, instead of the ASCAT3 allele-specific copy number data, because the previous integrative functional study that reported the TCGA-UVM data used ABSOLUTE^3^. We nevertheless checked that our results hold true even when ASCAT3 were used. All boxplots used either TCGA-UVM TPM expression data or ABSOLUTE copy number. SciPy’s (v1.16.0) ranksums function with default parameters was used to compute p-values in the boxplots grouped by *MITF* copy number (Figure 4C,D; 6E,F). Of the 80 TCGA-UVM patients, 37 had MITF copy = 1, 42 had MITF copy = 2, and 1 patient had MITF copy = 3. P-values shown in Figure 6A were calculated using the t-test (SciPy v1.16.0; ttest_ind; unequal variances), which was preferred in this case due to small sample sizes: 3, 34, 17, and 25, respectively for the groups MITF = 1, MYC = 2; MITF = 1, MYC ≥ 3; MITF = 2, MYC = 2; and MITF = 2, MYC ≥ 3. In the comparison of MITF-regulated pigment gene expressions between M3 and D3 groups (Figure S4C), the ranksums function was again used to compute p-values, but these were further subjected to genome-wide p-value adjustment using R’s p.adjust function with false discovery rate (FDR) method.

### Analysis of MITF correlation with MYC:MAX targets

The Genomic Regions Enrichment of Annotations Tool^98^ (GREAT, v4.0.4) was used to determine MYC:MAX targets by using MAX ChIP-seq data in human primary melanocytes^48^. A gene was assigned to be a MYC:MAX target if a MAX ChIP-seq peak resided within 500bp upstream of its TSS. An identical procedure using MITF ChIP-seq in human primary melanocytes^45^ was performed for the determination of MITF targets. Genes on chr3 were classified as either being a MYC:MAX target gene or a background (non-target) gene. MITF target genes on chr3 were excluded from this analysis due to the possibility of shared binding sites between MYC:MAX and MITF, both of which could recognize E-box elements. The Spearman correlations between MITF expression and MYC:MAX target gene expressions (TPM) were calculated using TCGA-UVM patients with the MITF copy = 1 (Figure S5A, 37 patients) or 2 (Figure S5B, 42 patients). Genes which were not expressed (TPM = 0.0) in patients with MITF copy = 1 or 2 were excluded from their respective analyses. The statistical significance between the MYC:MAX target distribution and background distribution was computed using the Kolmogorov-Smirnov test implemented in SciPy stats (v1.16.0; ks_2samp; default parameters).

### Analysis of *ZBTB17* expression and deletion in the contexts of *MITF* deletion and *MYC*amplification

The relationship between *ZBTB17*, *MITF*, and *MYC* was explored using both TCGA-UVM and TCGA-SKCM expression data by computing the Spearman correlation between *ZBTB17* and *MITF* expression (TPM). Using ABSOLUTE LiftOver copy number, we found *ZBTB17* was preferentially co-deleted with *MITF* in TCGA-UVM patients by using a one-sided binomial test to compute the statistical significance of *ZBTB17* deletion probability when *MITF* copy = 1 (37 patients) relative to *MITF* copy ≥ 2 (43 patients). Similarly, the statistical significance of co-occurring *ZBTB17* deletion and *MYC* amplification was assessed by using a one-sided binomial test to compare *ZBTB17* deletion probability between *MYC* copy= 2 (20 patients) and *MYC* copy ≥ 3 (60 patients). SciPy’s (v1.16.0) binomtest was used in both cases.

### Bivariate regression analysis of MITF expression using MYC and HUWE1

Utilizing gene expression (TPM) data for MITF, MYC, and HUWE1 in the cohort of 80 TCGA-UVM patients, we performed bivariate regression analysis (Python, statsmodels (v0.14.5); OLS; default parameters) to predict the expression of MITF given MYC and HUWE1 as predictors. Both response (MITF) and predictor (MYC, HUWE1) variables were z-score standardized prior to fitting but transformed back afterwards for visualization. Linear regression was done without a y-intercept, because the data were first mean centered.

### Bimodality analysis for MITF, MYC, and GNAQ/11

We stratified the TCGA-UVM data and identified 34 patients with MITF copy number = 1 and *GNAQ*/*11* mutations. We used the ABSOLUTE copy number data, and the *GNAQ*/*11* mutation data were retrieved through UCSC Xena Browser^85^. We calculated the bimodality index^56^ for MITF, MYC, and GNAQ/11 for these patients. The bimodality index shows the degree to which a 1D distribution can be described as a two-component mixture model and has a recommended value of 1.1 as a cutoff (higher values are deemed bimodal). The R (v4.4.3) implementation, BimodalIndex (v1.1.11), was used to calculate the index and confirm the bimodal distributions of MITF, MYC and GNAQ/11 (Figure S5F).

### Estimating the frequency of uveal melanoma patients with M3 and 8q+

We integrated the copy number data from TCGA-UVM (ABSOLUTE LiftOver) and the information from Johansson et al.^2^ to estimate the frequency of the M3/8q+ UVM subgroup. Of the 80 TCGA-UVM patients, 37 (46.3%) carried M3, of which 34/37 (91.9%) carried 8q+. M3 and 8q+ status were determined using modal copy number. The Johannson at al. UVM patient data reported 103 patients, 59 of which carried M3 (57.3%) and 46/59 (78.0%) had “gain” or “high gain” in chr8q ploidy. Combined, there were 96/183 (52.5%) of patients with M3, and 80/96 (83.3%) of those patients carried 8q+.

### Survival analysis

Kaplan-Meier curves and log-rank tests were computed with Python’s lifelines^99^ package (v0.30.0). Overall survival data for Figure 7A were retrieved from TCGA-UVM. Relapse-free survival from TCGA-UVM for Figure 7B and Figure S6A was calculated by taking the time to first occurrence of either death or recurrence, which does not include instances of a second primary tumor. *MITF* and *MYC* ABSOLUTE copy number were used to determine chr3 and chr8q ploidy for TCGA-UVM data. Figure 7B and Figure S6A also included data from Johansson *et al*. that was labeled as category 3 or 4 (LOH in chr3 with WT copy number chr 8 or 8q+, respectively), with Figure 7B only including patients with M3, marked by having hemizygous chr3p ploidy. Patients with status ‘Unknown’ were removed from the analysis.

### DepMap CRISPR screenings

Genome-wide CRISPR KO screening data for eight UVM cell lines (MEL202, MEL270, MEL285, MEL290, OMM25, UPMD1, UPMM3, WM3772F) were downloaded from DepMap^24^ (DepMap Public 25Q2+Score, Chronos; Figure 7C-F and Figure S6B-F). We calculated chr3 arithmetic mean expression from expression data available on “DepMap (Expression Public 25Q2),” utilizing all known genes on chr3 from the hg38 GENCODE V48 tracks (retrieved through UCSC Genome Browser Table Browser). Data for Figure S3B were taken from *CRISPR (DepMap Public 25Q2+Score, Chronos)*. WM3772F had no validated chr3 karyotyping data; we thus used chr3 expression analysis as a proxy for chr3 status.

## REFERENCES

1 Singh, A. D., Turell, M. E. & Topham, A. K. Uveal melanoma: trends in incidence, treatment, and survival. Ophthalmology 118, 1881–1885 (2011). 10.1016/j.ophtha.2011.01.040

2 Johansson, P. A. et al. Whole genome landscapes of uveal melanoma show an ultraviolet radiation signature in iris tumours. Nature communications 11, 2408 (2020). 10.1038/s41467-020-16276-8

3 Robertson, A. G. et al. Integrative Analysis Identifies Four Molecular and Clinical Subsets in Uveal Melanoma. Cancer Cell 32, 204–220 e215 (2017). 10.1016/j.ccell.2017.07.003

4 Prescher, G. et al. Prognostic implications of monosomy 3 in uveal melanoma. Lancet 347, 1222–1225 (1996). 10.1016/s0140-6736(96)90736-9

5 Parrella, P., Sidransky, D. & Merbs, S. L. Allelotype of posterior uveal melanoma: implications for a bifurcated tumor progression pathway. Cancer Res 59, 3032–3037 (1999).

6 Harbour, J. W. et al. Frequent mutation of BAP1 in metastasizing uveal melanomas. Science 330, 1410–1413 (2010). 10.1126/science.1194472

7 van de Nes, J. A. et al. Comparing the Prognostic Value of BAP1 Mutation Pattern, Chromosome 3 Status, and BAP1 Immunohistochemistry in Uveal Melanoma. Am J Surg Pathol 40, 796–805 (2016). 10.1097/PAS.0000000000000645

8 Durante, M. A. et al. Single-cell analysis reveals new evolutionary complexity in uveal melanoma. Nature communications 11, 496 (2020). 10.1038/s41467-019-14256-1

9 Baker, T. M., Waise, S., Tarabichi, M. & Van Loo, P. Aneuploidy and complex genomic rearrangements in cancer evolution. Nat Cancer 5, 228–239 (2024). 10.1038/s43018-023-00711-y

10 Gao, R. et al. Punctuated copy number evolution and clonal stasis in triple-negative breast cancer. Nat Genet 48, 1119–1130 (2016). 10.1038/ng.3641

11 Park, I. Y. et al. Dual Chromatin and Cytoskeletal Remodeling by SETD2. Cell 166, 950–962 (2016). 10.1016/j.cell.2016.07.005

12 Chiang, Y. C. et al. SETD2 Haploinsufficiency for Microtubule Methylation Is an Early Driver of Genomic Instability in Renal Cell Carcinoma. Cancer Res 78, 3135–3146 (2018). 10.1158/0008-5472.CAN-17-3460

13 Sharda, A. & Humphrey, T. C. The role of histone H3K36me3 writers, readers and erasers in maintaining genome stability. DNA Repair (Amst) 119, 103407 (2022). 10.1016/j.dnarep.2022.103407

14 Mason, F. M. et al. SETD2 safeguards the genome against isochromosome formation. Proc Natl Acad Sci U S A 120, e2303752120 (2023). 10.1073/pnas.2303752120

15 Levine, M. S. & Holland, A. J. The impact of mitotic errors on cell proliferation and tumorigenesis. Genes Dev 32, 620–638 (2018). 10.1101/gad.314351.118

16 Gisselsson, D. et al. Chromosomal breakage-fusion-bridge events cause genetic intratumor heterogeneity. Proc Natl Acad Sci U S A 97, 5357–5362 (2000). 10.1073/pnas.090013497

17 Khan, A. et al. A SETD2-CDK1-lamin axis maintains nuclear morphology and genome stability. Nat Cell Biol 27, 1327–1341 (2025). 10.1038/s41556-025-01723-9

18. Grady, D. Fragmentation of Rings and Shells. The Legacy of N.F. Mott., (Springer-Verlag, 2006).

19 Grady, D. E. Length scales and size distributions in dynamic fragmentation. International Journal of Fracture 163, 85–99 (2010).

20 Chen, X. et al. RasGRP3 Mediates MAPK Pathway Activation in GNAQ Mutant Uveal Melanoma. Cancer Cell 31, 685–696 e686 (2017). 10.1016/j.ccell.2017.04.002

21 Elbatsh, A. M. O. et al. INPP5A phosphatase is a synthetic lethal target in GNAQ and GNA11-mutant melanomas. Nat Cancer 5, 481–499 (2024). 10.1038/s43018-023-00710-z

22 Chan, P. Y. et al. The synthetic lethal interaction between CDS1 and CDS2 is a vulnerability in uveal melanoma and across multiple tumor types. Nat Genet 57, 1672–1683 (2025). 10.1038/s41588-025-02222-1

23 Levy, C., Khaled, M. & Fisher, D. E. MITF: master regulator of melanocyte development and melanoma oncogene. Trends Mol Med 12, 406–414 (2006). 10.1016/j.molmed.2006.07.008

24 Arafeh, R., Shibue, T., Dempster, J. M., Hahn, W. C. & Vazquez, F. The present and future of the Cancer Dependency Map. Nat Rev Cancer 25, 59–73 (2025). 10.1038/s41568-024-00763-x

25 Maitra, A. et al. High-resolution chromosome 3p allelotyping of breast carcinomas and precursor lesions demonstrates frequent loss of heterozygosity and a discontinuous pattern of allele loss. Am J Pathol 159, 119–130 (2001). 10.1016/S0002-9440(10)61679-3

26. Cancer Genome Atlas Research, N. Comprehensive molecular characterization of clear cell renal cell carcinoma. Nature 499, 43-49 (2013). 10.1038/nature12222

27. Cancer Genome Atlas Research, N., et al. Comprehensive Molecular Characterization of Papillary Renal-Cell Carcinoma. The New England journal of medicine 374, 135-145 (2016). 10.1056/NEJMoa1505917

28 Shridhar, R. et al. Frequent breakpoints in the 3p14.2 fragile site, FRA3B, in pancreatic tumors. Cancer Res 56, 4347–4350 (1996).

29 Yagyu, T. et al. Human chromosome 3p21.3 carries TERT transcriptional regulators in pancreatic cancer. Sci Rep 11, 15355 (2021). 10.1038/s41598-021-94711-6

30 Garraway, L. A. et al. Integrative genomic analyses identify MITF as a lineage survival oncogene amplified in malignant melanoma. Nature 436, 117–122 (2005). 10.1038/nature03664

31 Umbreit, N. T. et al. Mechanisms generating cancer genome complexity from a single cell division error. Science 368 (2020). 10.1126/science.aba0712

32 Stephens, P. J. et al. Massive genomic rearrangement acquired in a single catastrophic event during cancer development. Cell 144, 27–40 (2011). 10.1016/j.cell.2010.11.055

33 Vitale, I., Galluzzi, L., Castedo, M. & Kroemer, G. Mitotic catastrophe: a mechanism for avoiding genomic instability. Nat Rev Mol Cell Biol 12, 385–392 (2011). 10.1038/nrm3115

34 Aymard, F. et al. Transcriptionally active chromatin recruits homologous recombination at DNA double-strand breaks. Nat Struct Mol Biol 21, 366–374 (2014). 10.1038/nsmb.2796

35 Liu, P. et al. Chromosome catastrophes involve replication mechanisms generating complex genomic rearrangements. Cell 146, 889–903 (2011). 10.1016/j.cell.2011.07.042

36 Lundberg, A. et al. The long-term prognostic and predictive capacity of cyclin D1 gene amplification in 2305 breast tumours. Breast Cancer Res 21, 34 (2019). 10.1186/s13058-019-1121-4

37 Bocanegra, M. et al. Focal amplification and oncogene dependency of GAB2 in breast cancer. Oncogene 29, 774–779 (2010). 10.1038/onc.2009.364

38 Dilshat, R. et al. MITF reprograms the extracellular matrix and focal adhesion in melanoma. eLife 10 (2021). 10.7554/eLife.63093

39 Tong, X. et al. Crosstalk in Skin: Loss of Desmoglein 1 in Keratinocytes Inhibits BRAF(V600E)-Induced Cellular Senescence in Human Melanocytes. J Invest Dermatol 145, 1740–1752 e1744 (2025). 10.1016/j.jid.2024.10.608

40 Chen, Y. et al. Co-isolation of human donor eye cells and development of oncogene-mutated melanocytes to study uveal melanoma. BMC Biol 23, 16 (2025). 10.1186/s12915-025-02118-w

41 Sun, P., Enslen, H., Myung, P. S. & Maurer, R. A. Differential activation of CREB by Ca2+/calmodulin-dependent protein kinases type II and type IV involves phosphorylation of a site that negatively regulates activity. Genes Dev 8, 2527–2539 (1994). 10.1101/gad.8.21.2527

42 Melville, Z. et al. The Activation of Protein Kinase A by the Calcium-Binding Protein S100A1 Is Independent of Cyclic AMP. Biochemistry 56, 2328–2337 (2017). 10.1021/acs.biochem.7b00117

43 Motiani, R. K. et al. STIM1 activation of adenylyl cyclase 6 connects Ca(2+) and cAMP signaling during melanogenesis. EMBO J 37 (2018). 10.15252/embj.201797597

44 Seoane, M. et al. Lineage-specific control of TFIIH by MITF determines transcriptional homeostasis and DNA repair. Oncogene 38, 3616–3635 (2019). 10.1038/s41388-018-0661-x

45 Webster, D. E. et al. Enhancer-targeted genome editing selectively blocks innate resistance to oncokinase inhibition. Genome Res 24, 751–760 (2014). 10.1101/gr.166231.113

46 Cho, S. W. et al. Promoter of lncRNA Gene PVT1 Is a Tumor-Suppressor DNA Boundary Element. Cell 173, 1398–1412 e1322 (2018). 10.1016/j.cell.2018.03.068

47 Tseng, Y. Y. et al. PVT1 dependence in cancer with MYC copy-number increase. Nature 512, 82–86 (2014). 10.1038/nature13311

48 Hejna, M. et al. Local genomic features predict the distinct and overlapping binding patterns of the bHLH-Zip family oncoproteins MITF and MYC-MAX. Pigment Cell Melanoma Res 32, 500–509 (2019). 10.1111/pcmr.12762

49 Luo, Y. et al. New developments on the Encyclopedia of DNA Elements (ENCODE) data portal. Nucleic Acids Res 48, D882–D889 (2020). 10.1093/nar/gkz1062

50 Soshnikova, N. V. et al. PHF10 subunit of PBAF complex mediates transcriptional activation by MYC. Oncogene 40, 6071–6080 (2021). 10.1038/s41388-021-01994-0

51 Zacarias-Fluck, M. F. et al. Reducing MYC’s transcriptional footprint unveils a good prognostic gene signature in melanoma. Genes Dev 37, 303–320 (2023). 10.1101/gad.350078.122

52 Walz, S. et al. Activation and repression by oncogenic MYC shape tumour-specific gene expression profiles. Nature 511, 483–487 (2014). 10.1038/nature13473

53 Peter, S. et al. Tumor cell-specific inhibition of MYC function using small molecule inhibitors of the HUWE1 ubiquitin ligase. EMBO molecular medicine 6, 1525–1541 (2014). 10.15252/emmm.201403927

54 Venturelli, O. S., El-Samad, H. & Murray, R. M. Synergistic dual positive feedback loops established by molecular sequestration generate robust bimodal response. Proc Natl Acad Sci U S A 109, E3324–3333 (2012). 10.1073/pnas.1211902109

55 Shi, C., Li, H. X. & Zhou, T. Architecture-dependent robustness in a class of multiple positive feedback loops. IET Syst Biol 7, 1–10 (2013). 10.1049/iet-syb.2011.0090

56 Wang, J., Wen, S., Symmans, W. F., Pusztai, L. & Coombes, K. R. The bimodality index: a criterion for discovering and ranking bimodal signatures from cancer gene expression profiling data. Cancer Inform 7, 199–216 (2009). 10.4137/cin.s2846

57 Carreira, S. et al. Mitf regulation of Dia1 controls melanoma proliferation and invasiveness. Genes Dev 20, 3426–3439 (2006). 10.1101/gad.406406

58 Hoek, K. S. & Goding, C. R. Cancer stem cells versus phenotype-switching in melanoma. Pigment Cell Melanoma Res 23, 746–759 (2010). 10.1111/j.1755-148X.2010.00757.x

59 Piaggio, F. et al. In uveal melanoma Galpha-protein GNA11 mutations convey a shorter disease-specific survival and are more strongly associated with loss of BAP1 and chromosomal alterations than Galpha-protein GNAQ mutations. Eur J Cancer 170, 27–41 (2022). 10.1016/j.ejca.2022.04.013

60 Nareyeck, G., Zeschnigk, M., Bornfeld, N. & Anastassiou, G. Novel cell lines derived by long-term culture of primary uveal melanomas. Ophthalmologica 223, 196–201 (2009). 10.1159/000201566

61 Mergener, S., Siveke, J. T. & Pena-Llopis, S. Monosomy 3 Is Linked to Resistance to MEK Inhibitors in Uveal Melanoma. Int J Mol Sci 22 (2021). 10.3390/ijms22136727

62 Liu, H. et al. The Chick Chorioallantoic Membrane as a Xenograft Model for the Quantitative Analysis of Uveal Melanoma Metastasis in Multiple Organs. Cells 13 (2024). 10.3390/cells13141169

63. The Wistar Institute Melanoma Research Center, <https://www.wistar.org/our-scientists/meenhard-herlyn> (

64. Jager, M. J., Magner, J. A., Ksander, B. R. & Dubovy, S. R. Uveal Melanoma Cell Lines: Where do they come from? (An American Ophthalmological Society Thesis). Trans Am Ophthalmol Soc 114, T5 (2016).

65 Ferrier, S. T., Li, M. & Burnier, J. V. Azacytidine treatment affects the methylation pattern of genomic and cell-free DNA in uveal melanoma cell lines. BMC Cancer 24, 1299 (2024). 10.1186/s12885-024-13037-4

66 Nareyeck, G., Zeschnigk, M., Prescher, G., Lohmann, D. R. & Anastassiou, G. Establishment and characterization of two uveal melanoma cell lines derived from tumors with loss of one chromosome 3. Exp Eye Res 83, 858–864 (2006). 10.1016/j.exer.2006.04.004

67 Herwig-Carl, M. C. et al. Mass Spectrometry-Based Profiling of Histone Post-Translational Modifications in Uveal Melanoma Tissues, Human Melanocytes, and Uveal Melanoma Cell Lines - A Pilot Study. Invest Ophthalmol Vis Sci 65, 27 (2024). 10.1167/iovs.65.2.27

68 Goldrick, C. et al. Hindsight: Review of Preclinical Disease Models for the Development of New Treatments for Uveal Melanoma. J Cancer 12, 4672–4685 (2021). 10.7150/jca.53954

69 Angeloni, D. Molecular analysis of deletions in human chromosome 3p21 and the role of resident cancer genes in disease. Brief Funct Genomic Proteomic 6, 19–39 (2007). 10.1093/bfgp/elm007

70 Lovrecic, L., Bertok, S. & Zerjav Tansek, M. A New Case of an Extremely Rare 3p21.31 Interstitial Deletion. Mol Syndromol 7, 93–98 (2016). 10.1159/000445227

71 Malmgren, H., Sahlen, S., Wide, K., Lundvall, M. & Blennow, E. Distal 3p deletion syndrome: detailed molecular cytogenetic and clinical characterization of three small distal deletions and review. Am J Med Genet A **143A**, 2143–2149 (2007). 10.1002/ajmg.a.31902

72 Brien, G. L. et al. Polycomb PHF19 binds H3K36me3 and recruits PRC2 and demethylase NO66 to embryonic stem cell genes during differentiation. Nat Struct Mol Biol 19, 1273–1281 (2012). 10.1038/nsmb.2449

73 Mertens, F., Johansson, B. & Mitelman, F. Isochromosomes in neoplasia. Genes Chromosomes Cancer 10, 221–230 (1994). 10.1002/gcc.2870100402

74 Izumi, K. & Krantz, I. D. Pallister-Killian syndrome. Am J Med Genet C Semin Med Genet 166C, 406–413 (2014). 10.1002/ajmg.c.31423

75 Ibarra-Ramirez, M., Campos-Acevedo, L. D. & Martinez de Villarreal, L. E. Chromosomal Abnormalities of Interest in Turner Syndrome: An Update. J Pediatr Genet 12, 263–272 (2023). 10.1055/s-0043-1770982

76 Bolkestein, M. et al. Chromothripsis in Human Breast Cancer. Cancer Res 80, 4918–4931 (2020). 10.1158/0008-5472.CAN-20-1920

77 Maresca, V. et al. Linking alphaMSH with PPARgamma in B16-F10 melanoma. Pigment Cell Melanoma Res 26, 113–127 (2013). 10.1111/j.1755-148X.2012.01042.x

78 Tanwar, J. et al. MITF is a novel transcriptional regulator of the calcium sensor STIM1: Significance in physiological melanogenesis. J Biol Chem 298, 102681 (2022). 10.1016/j.jbc.2022.102681

79 Shaughnessy, M. et al. Selective uveal melanoma inhibition with calcium channel blockade. Int J Oncol 55, 1090–1096 (2019). 10.3892/ijo.2019.4873

80 Van Raamsdonk, C. D., Fitch, K. R., Fuchs, H., de Angelis, M. H. & Barsh, G. S. Effects of G-protein mutations on skin color. Nat Genet 36, 961–968 (2004). 10.1038/ng1412

81 Aida, S. et al. MITF suppression improves the sensitivity of melanoma cells to a BRAF inhibitor. Cancer Lett 409, 116–124 (2017). 10.1016/j.canlet.2017.09.008

82 Llombart, V. & Mansour, M. R. Therapeutic targeting of "undruggable" MYC. EBioMedicine 75, 103756 (2022). 10.1016/j.ebiom.2021.103756

83 Garralda, E. et al. MYC targeting by OMO-103 in solid tumors: a phase 1 trial. Nat Med 30, 762–771 (2024). 10.1038/s41591-024-02805-1

84 Singleton, K. R. et al. Melanoma Therapeutic Strategies that Select against Resistance by Exploiting MYC-Driven Evolutionary Convergence. Cell reports 21, 2796–2812 (2017). 10.1016/j.celrep.2017.11.022

85 Goldman, M. J. et al. Visualizing and interpreting cancer genomics data via the Xena platform. Nat Biotechnol 38, 675–678 (2020). 10.1038/s41587-020-0546-8

86 Perez, G. et al. The UCSC Genome Browser database: 2025 update. Nucleic Acids Res 53, D1243–D1249 (2025). 10.1093/nar/gkae974

87 Love, M. I., Huber, W. & Anders, S. Moderated estimation of fold change and dispersion for RNA-seq data with DESeq2. Genome Biol 15, 550 (2014). 10.1186/s13059-014-0550-8

88 Pimentel, H., Bray, N. L., Puente, S., Melsted, P. & Pachter, L. Differential analysis of RNA-seq incorporating quantification uncertainty. Nat Methods 14, 687–690 (2017). 10.1038/nmeth.4324

89 Corrales, E. et al. Dynamic transcriptome analysis reveals signatures of paradoxical effect of vemurafenib on human dermal fibroblasts. Cell Commun Signal 19, 123 (2021). 10.1186/s12964-021-00801-3

90 Smith, L. K. et al. Adaptive translational reprogramming of metabolism limits the response to targeted therapy in BRAF(V600) melanoma. Nature communications 13, 1100 (2022). 10.1038/s41467-022-28705-x

91 Peng, J. et al. Vemurafenib induces a noncanonical senescence-associated secretory phenotype in melanoma cells which promotes vemurafenib resistance. Heliyon 9, e17714 (2023). 10.1016/j.heliyon.2023.e17714

92 Flower, C. T. et al. Signaling and transcriptional dynamics underlying early adaptation to oncogenic BRAF inhibition. bioRxiv (2024). 10.1101/2024.02.19.581004

93 Kim, D., Paggi, J. M., Park, C., Bennett, C. & Salzberg, S. L. Graph-based genome alignment and genotyping with HISAT2 and HISAT-genotype. Nat Biotechnol 37, 907–915 (2019). 10.1038/s41587-019-0201-4

94 Shumate, A., Wong, B., Pertea, G. & Pertea, M. Improved transcriptome assembly using a hybrid of long and short reads with StringTie. PLoS Comput Biol 18, e1009730 (2022). 10.1371/journal.pcbi.1009730

95 Wen, J. et al. Modeling of pigmentation disorders associated with MITF mutation in Waardenburg syndrome revealed an impaired melanogenesis pathway in iPS-derived melanocytes. Pigment Cell Melanoma Res 37, 21–35 (2024). 10.1111/pcmr.13118

96. Johns, E. et al. The Lipid Droplet Protein DHRS3 Is a Regulator of Melanoma Cell State. Pigment Cell Melanoma Res 38, e13208 (2025). 10.1111/pcmr.13208

97 Ritchie, M. E. et al. limma powers differential expression analyses for RNA-sequencing and microarray studies. Nucleic Acids Res 43, e47 (2015). 10.1093/nar/gkv007

98 McLean, C. Y. et al. GREAT improves functional interpretation of cis-regulatory regions. Nat Biotechnol 28, 495–501 (2010). 10.1038/nbt.1630

99 Davidson-Pilon, C. lifelines: survival analysis in Python. Journal of Open Source Software 4, 1317 (2019). 10.21105/joss.01317

