## Supplementary Figures for "Chromosome 3p deletion leads to extensive genomic alterations in diverse cancers and confers synthetic lethality in uveal melanoma"

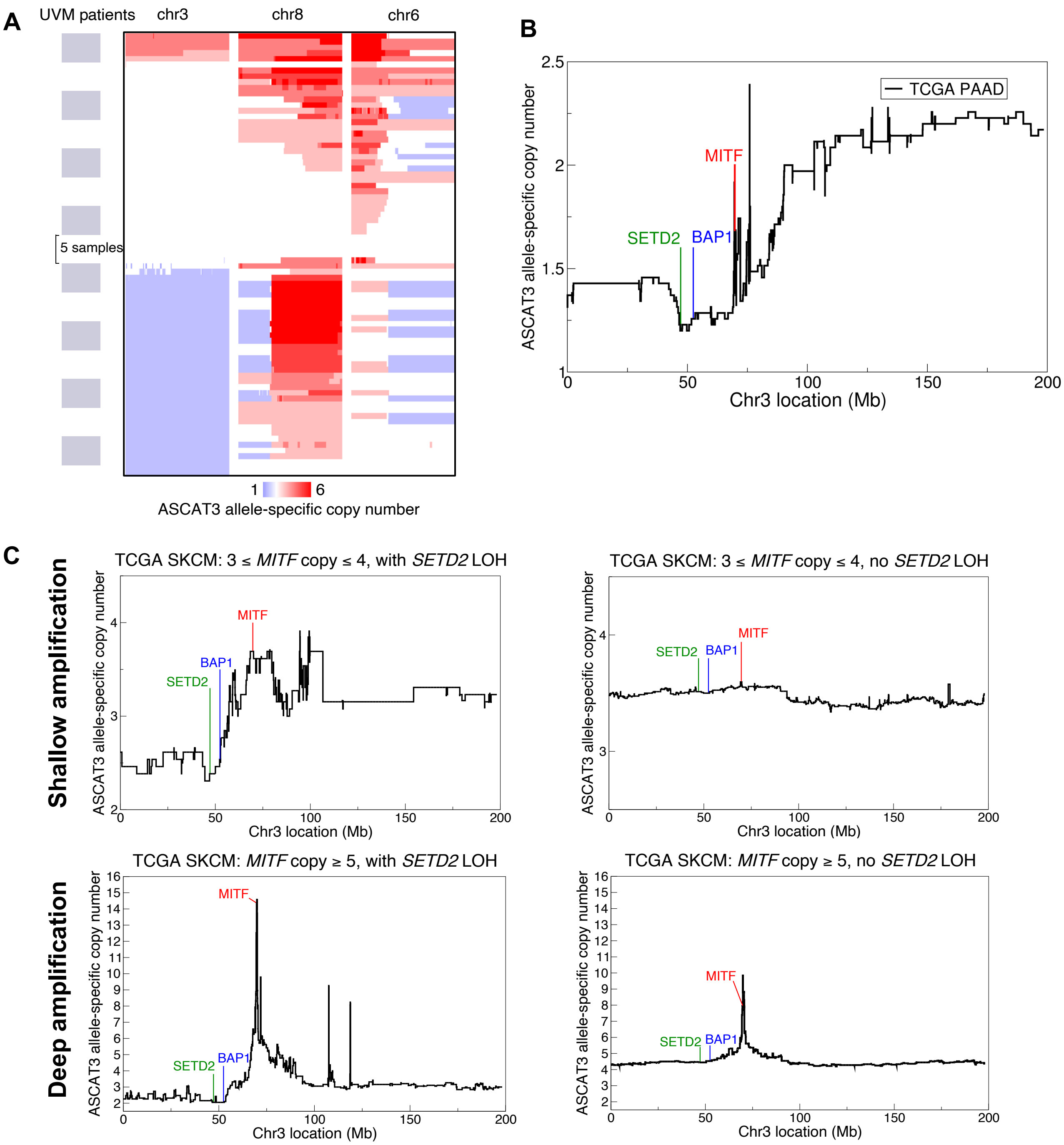


**Figure S1. Deletion patterns of chr3 in UVM and SKCM, related to Figure 1.** (A) Chr3, chr6, and chr8 allele-specific copy number data for 77 TCGA UVM patients. Copy number loss (blue) and gain (red) reveals isochromosome formation on chr6 and chr8 concurrent with chr3 loss. The figure was modified from a Xena Browser (85) screen shot. (B) Mean chr3 deletion profile in 35 TCGA PAAD patients. One patient had a deep amplification (copy=13) of *MITF* when *SETD2* and *BAP1* were deleted, creating the local spike in the figure. Otherwise, *MITF* was on the path of Poisson breaks. (C) Averaged patterns of shallow amplification of *MITF* with *SETD2* LOH (13 patients), shallow amplification of *MITF* without *SETD2* LOH (157 patients), deep amplification of *MITF* with *SETD2* LOH (19 patients), and deep amplification of *MITF* without *SETD2* LOH (36 patients) in TCGA SKCM data. Deep MITF amplification is seen to be more pronounced, roughly 2-folder higher, in the context of chr3p LOH.

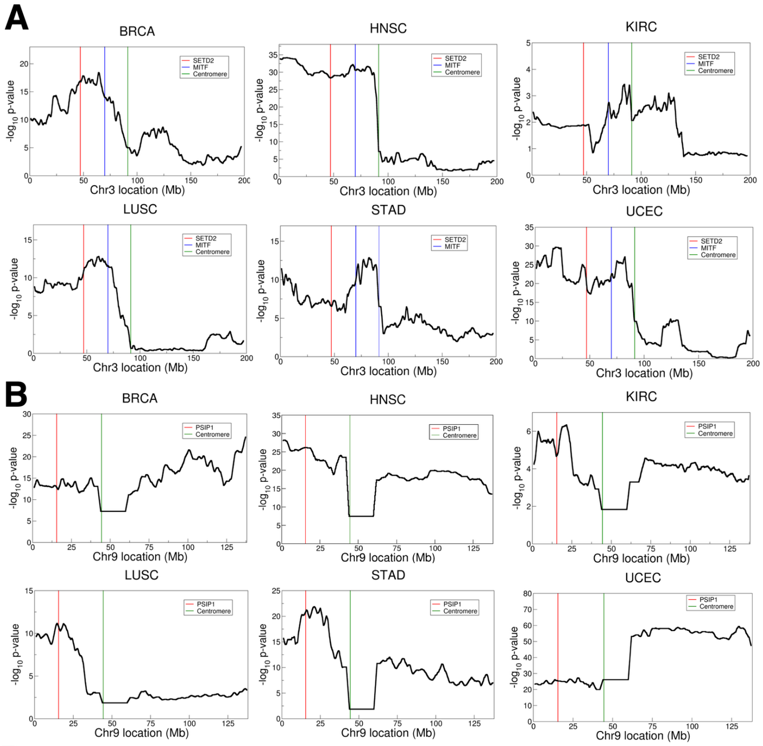


C

| Feature | Estimate | p-value |
|---|---|---|
| Intercept | 2.4 | $1.8 \times 10^{-6}$ |
| Distance | 0.08 | $< 2.0 \times 10^{-16}$ |
| CESC | -2.2 | $1.3 \times 10^{-5}$ |
| COAD | -2.0 | $5.1 \times 10^{-5}$ |
| HNSC | -1.9 | $9.3 \times 10^{-5}$ |
| KIRC | -2.5 | $1.5 \times 10^{-6}$ |
| LGG | -2.2 | $2.6 \times 10^{-5}$ |
| LIHC | -2.1 | $1.3 \times 10^{-5}$ |
| LUSC | -1.9 | $4.9 \times 10^{-5}$ |
| OV | -2.3 | $1.2 \times 10^{-6}$ |
| PCPG | -2.9 | $1.0 \times 10^{-5}$ |
| PRAD | -2.2 | $1.1 \times 10^{-5}$ |
| TGCT | -2.9 | $5.7 \times 10^{-6}$ |
| chr7 | 0.9 | $3.7 \times 10^{-5}$ |
| chr8 | 1.1 | $8.1 \times 10^{-11}$ |
| chr11 | 1.8 | $< 2.0 \times 10^{-16}$ |
| chr12 | 1.3 | $3.0 \times 10^{-10}$ |
| chr18 | 1.0 | $8.4 \times 10^{-7}$ |
| chr19 | 1.2 | $5.1 \times 10^{-9}$ |
| chr20 | 0.9 | $2.7 \times 10^{-5}$ |

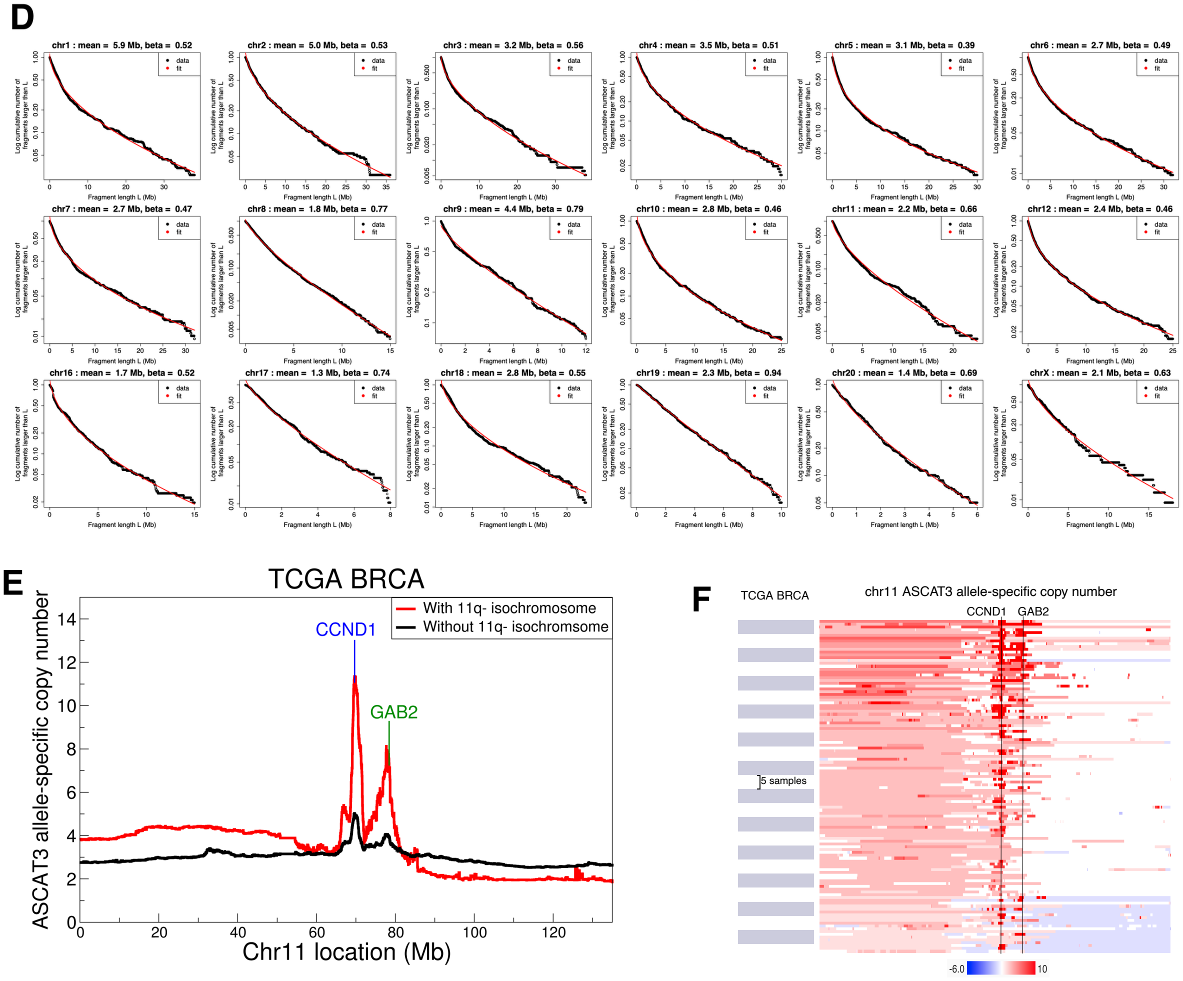

**Figure S2. *SETD2* and *PSIP1* loss are implicated in isochromosome formation and chromosome fragmentation in human cancers, related to Figure 2.** (A) Wald test p-value for observing higher isochromosome occurrence probability in patients with LOH at each chr3 location compared to patients without LOH at the same location. Chr3 locations were evenly sampled at 100kb intervals, and the transformed p-values were smoothed using a running window of 20 samples. (B) Same as in (A), but for chr9. (C) Significant coefficients ($p<10^{-4}$) in multivariate regression of number of heterozygous fragments on the LOH arm of 11,658 isochromosomes found in 33 TCGA cancers against fusion site distance, cancer type, chromosomes and intercept. (D) $Log_{10}$ cumulative distribution of fragment sizes for each chromosome, pooled across cancers. (E) Mean chr11q amplification profile in TCGA-BRCA patients with 11q- isochromosomes (red line, 117 patients) vs. without 11q- isochromosomes (black line, 580 patients). Only those patients with an amplification event on chr11q are shown. The mean copy numbers of the driver genes *CCND1* and *GAB2* on chr11q are ~2-fold higher in the patients with 11q- isochromosomes than those without 11q- isochromosomes. (F) Heatmap of chr11 ASCAT3 allele-specific copy number data for TCGA-BRCA patients with 11q- isochromosomes shown in (E). It can be seen that the *CCND1* and *GAB2* loci acquired deep amplifications via repeated fragmentation of the centromeric 11q segments on dicentric isochromosomes. The figure was modified from a Xena Browser (85) screen shot.

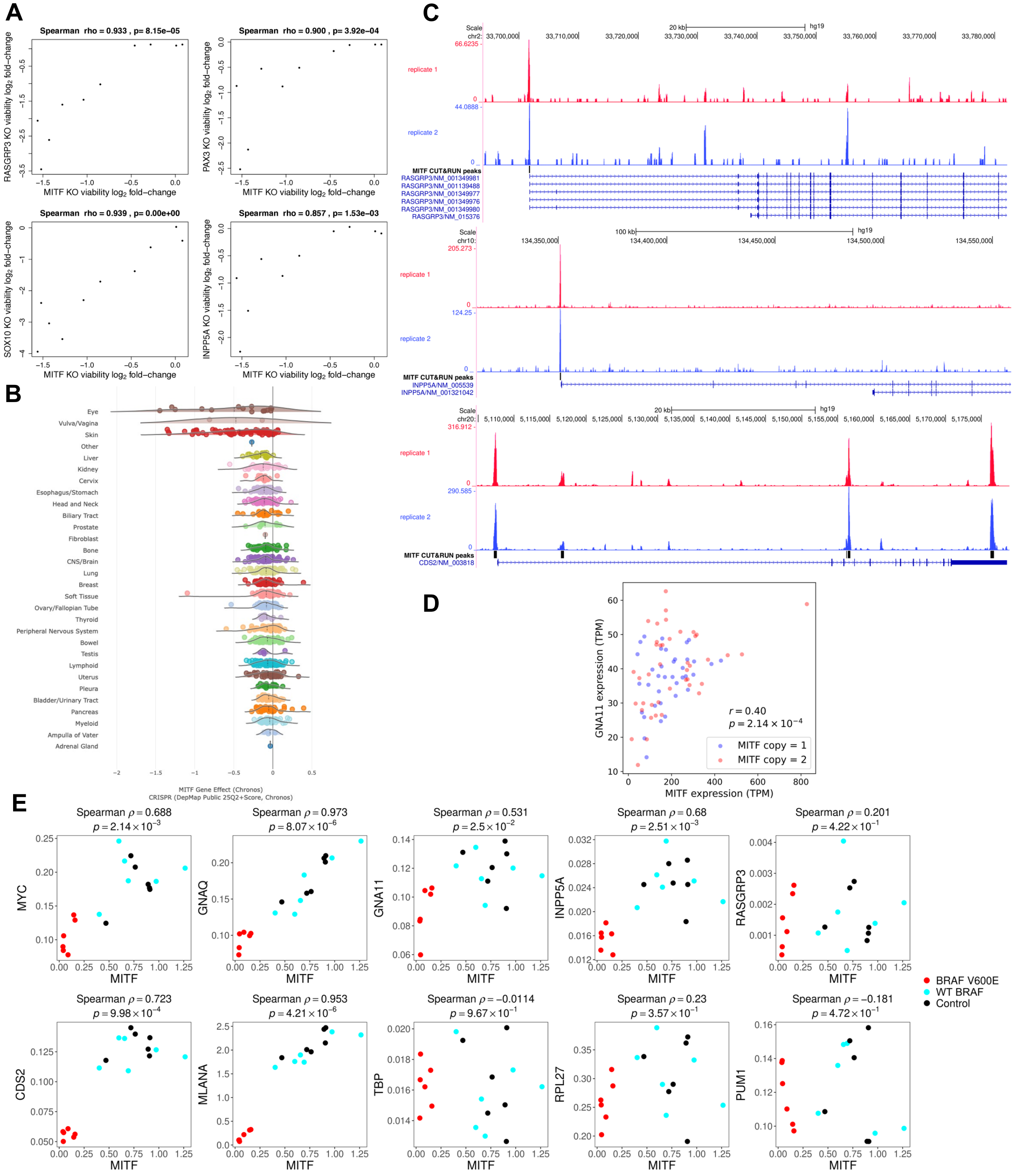


**Figure S3. MITF regulates several synthetic-lethal genes and is crucial for UVM viability, related to Figure 3.** (A) Elbatsh *et al.*'s CRISPR screening results in UVM cell lines, combining cell viability data from days 14 and 21 for MITF, RASGRP3, PAX3, SOX10, and INPP5A. Strong correlation is found between the KO viability $\log_2$ fold-change of MITF and its transcriptional targets RASGRP3 and INPP5A as well as the cooperating factors PAX3 and SOX10. (B) CRISPR KO gene effect of MITF on all available cell lines in DepMap, grouped by tissue type. MITF KO has the most significant effect on ocular melanoma (top row), even compared to skin cutaneous melanoma (third row). The category *eye* contains ocular melanoma, non-cancerous immortalized cells and retinoblastoma; the gene effect of MITF KO on non-cancerous cells and retinoblastoma was greater than -0.272. (C) MITF CUT&RUN (38) in SK-MEL-28 shows called peaks in the promoter of *RASA4B*, *INPP5A*, and *CDS2*. (D) GNA11 vs. MITF expression in TCGA-UVM data, colored by MITF copy number. Pearson correlation coefficient is calculated by pooling the M3 and D3 patients together. (E) Gene expression in neonatal foreskin melanocytes transduced with WT BRAF or $BRAF^{V600E}$ and treated with conditioned media from keratinocytes (39) (GEO GSE255546). Both shControl and shDsg1 data are used in the analysis. The gene-level counts were normalized by the total counts in each sample and then multiplied by 1000 for visualization. The data show correlated normalized gene expression for most MITF targets (*MYC*, *GNAQ*, *GNA11*, *INPP5A*, *RASGRP3*, *CDS2*, *MLANA*). The housekeeping genes *TBP*, *RPL27*, and *PUM1* were used as an additional control.

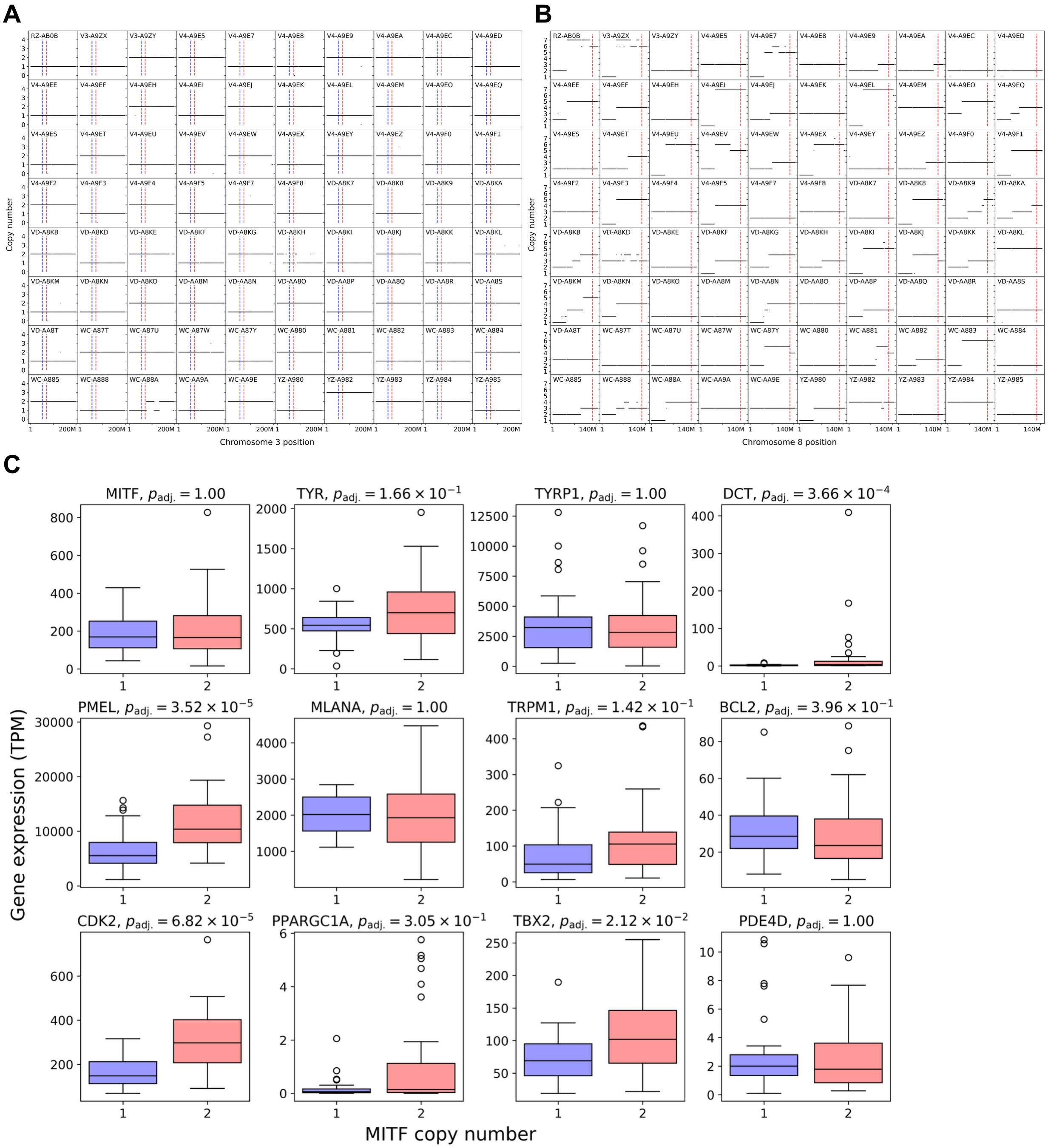


**Figure S4. Chr3 deletion and 8q amplification preferentially co-occur in UVM, leaving the expression of *MITF*-regulated pigment genes unchanged, related to Figure 4.** (A) ABSOLUTE LiftOver gene-level copy number for chr3 for the cohort of 80 TCGA-UVM patients. The blue and red lines indicate the locations of *BAP1* and *MITF*, respectively. Only the tissue source site and participant identifiers were retained in each TCGA barcode. (B) Same as (A), but on chr8 and with a red line indicating the position of *MYC*. (C) Boxplots of the expression levels of known MITF target genes for M3 (MITF copy=1) and D3 (MITF copy=2) TCGA-UVM patients. A Wilcoxon rank-sum test shows that most MITF-regulated pigment genes remain unchanged in expression between M3 and D3 patients. P-values were adjusted using genome-wide FDR.

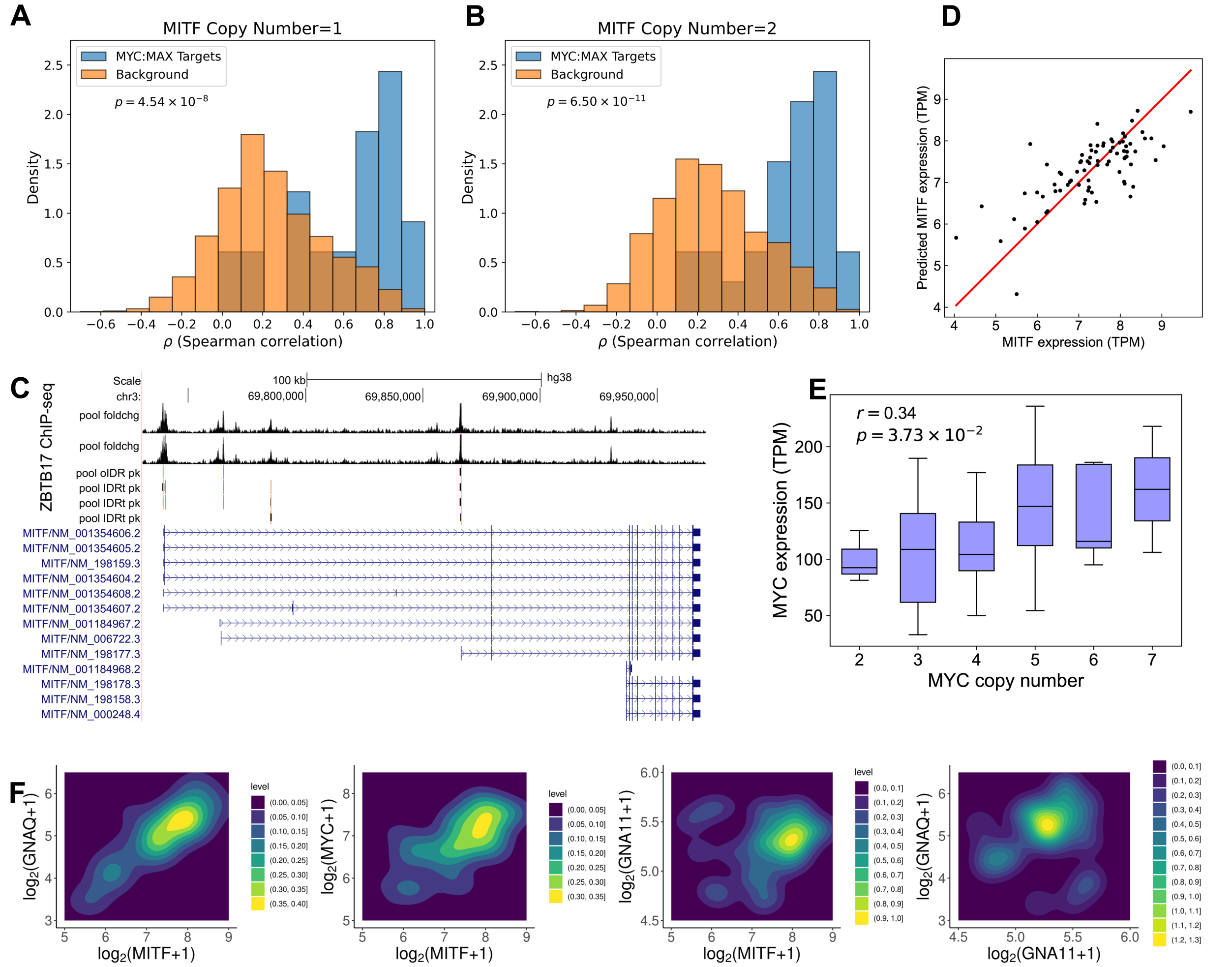


**Figure S5. MYC regulates MITF and helps create bistable states in M3 UVM, related to Figure 5 and Figure 6.** (A) The distribution of correlations between MITF expression and MYC:MAX target gene expression levels on chr3 (blue) compared to the background distribution of correlations between MITF and non-target genes of MYC:MAX on chr3 (orange) in M3 patients. The significant difference between the distributions (Kolmogorov-Smirnov test) shows that MITF strongly correlates with MYC:MAX targets. MYC:MAX-target genes were determined by assigning MAX ChIP-seq peaks (used as a MYC proxy) to a gene if the peak occurred within 500bp upstream of the TSS. MITF targets (45) were removed from both MYC:MAX-target and background distributions. (B) Same as (A), but for *MITF* copy number $= 2$. (C) ENCODE ZBTB17 ChIP-seq (49) in HEK-293 cell line indicates that ZBTB17 binds several locations within *MITF*. (D) Observed vs. predicted MITF expression in TCGA-UVM based on bivariate regression of *MITF* against *MYC* and *HUWE1* expression levels ($R^2 = 0.543$; MYC partial correlation = 0.584, $p$ = $3.2 \times 10^{-7}$; HUWE1 partial correlation = 0.24, $p$ = 0.024). (E) Boxplots of MYC expression levels grouped into different *MYC* copy numbers in the TCGA-UVM M3 cohort. MYC expression shows strong Pearson correlation with increasing MYC copy number when conditioned on M3. (F) Distributions of log GNAQ, MITF, MYC and GNA11 expression for 34 TCGA-UVM patients with M3 and GNAQ/11 mutations. Bistable low (GNAQ, MITF, MYC) and high (GNAQ, MITF, MYC) states are visible, consistent with the bimodality index being 1.2, 1.6, and 1.2 for GNAQ, MITF and MYC, respectively. The bimodality index for GNA11 was 0.96, indicating that a simple bimodal distribution was not a good fit. In addition to the low (GNAQ, MITF, GNA11) state, there was also a low (GNAQ, MITF) / high (GNA11) state that couldn't be explained by MITF-mediated regulation.

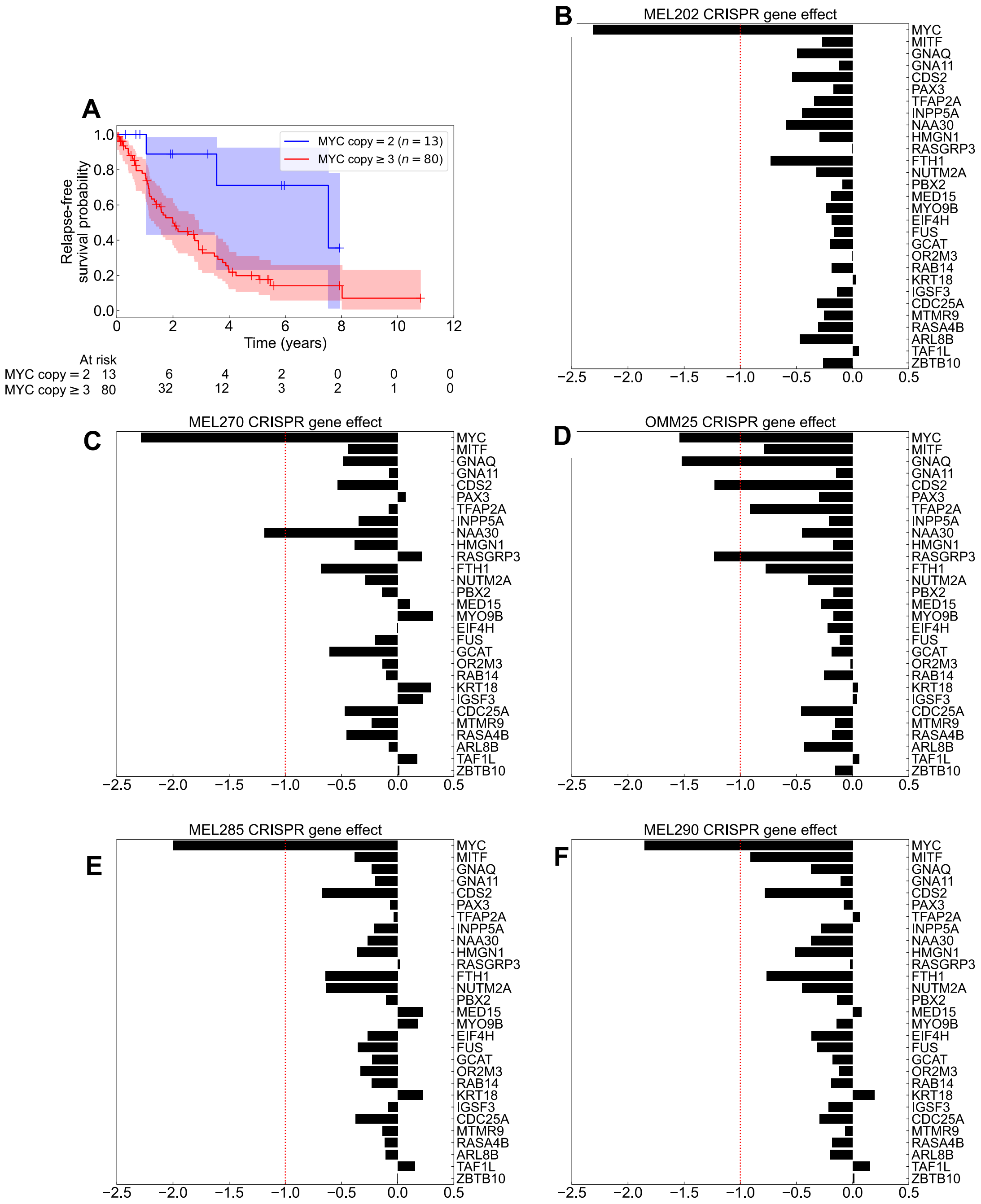


**Figure S6. Chr3 LOH with MYC amplification worsens prognosis and increases selective dependence of UVM on MITF, related to Figure 7.** (A) Kaplan-Meier survival curves for relapse-free survival of UVM patients with chr3 LOH, split into groups with *MYC* amplification or WT copy number. The shaded region indicates a 95% confidence interval; crosshairs mark right-censored data. (B-F) DepMap CRISPR gene effect scores in UVM cell lines with D3 for *GNAQ*/*GNA11*, synthetic-lethal genes identified by Elbatsh *et al.* (21) and the UVM-vulnerability gene *CDS2* identified by Chan *et al.* (22); genes with missing data are excluded. The dashed line at -1 is a recommended threshold for assessing high dependency (lower values). Apart from MYC, which is a common essential gene, MITF and most other genes do not pass the threshold in these D3 UVM. In sharp contrast, MITF KO has a pronounced effect on M3 UVM (Figure 7D,E). MEL202, MEL270, OMM25 are *GNAQ*-mutant. MEL285 and MEL290 have the wildtype *GNAQ/11*.
